# Shifts in mutation spectra enhance access to beneficial mutations

**DOI:** 10.1101/2020.09.05.284158

**Authors:** Mrudula Sane, Gaurav D Diwan, Bhoomika A Bhat, Lindi M Wahl, Deepa Agashe

## Abstract

Biased mutation spectra are pervasive, with wide variation in the magnitude of mutational biases that influence genome evolution and adaptation. How do such diverse biases evolve? Our experiments show that changing the mutation spectrum allows populations to sample previously under-sampled mutational space, including beneficial mutations. The resulting shift in the distribution of fitness effects is advantageous: beneficial mutation supply and beneficial pleiotropy both increase, while deleterious load reduces. More broadly, simulations indicate that reducing or reversing the direction of a long-term bias is always selectively favoured. Such changes in mutation bias can occur easily via altered function of DNA repair genes. A phylogenetic analysis shows that these genes are repeatedly gained and lost in bacterial lineages, leading to frequent bias shifts in opposite directions. Thus, shifts in mutation spectra may evolve under selection, and can directly alter the outcome of adaptive evolution by facilitating access to beneficial mutations.

**SIGNIFICANCE STATEMENT:** Mutations are important because they provide raw material for evolution. Some types of mutations occur more often than others, and the strength of such mutational bias varies across species. It is not clear how this variation arises. We experimentally measured the immediate effects of changing the mutation bias of *E. coli*, and used simulations to understand the long-term effects. Altering mutational bias is beneficial whenever the new bias increases sampling of mutational classes that were previously under-sampled. We also show that historically, bacteria have often experienced such beneficial bias switches. Our work thus demonstrates the importance of mutational biases in evolution. By allowing exploration of new mutational space, altered mutation biases could drive rapid adaptation.

## INTRODUCTION

The mutation spectrum describes the frequency of various classes of sampled mutations (e.g., transversions (Tv) vs. transitions (Ts)), and is often governed by the action of DNA repair enzymes. By determining the pool of genetic variants available for selection, mutation spectra can shape key genome features (e.g., nucleotide (1, 2), codon and amino acid composition (3)), even if under selection for thousands of generations (4); determine the genetic basis of adaptation (5–9), driving convergent evolution (10–12); and shape the evolution of resistance to antibiotics (13–15) and anti-cancer drugs (16). Thus, mutation spectra are important for adaptive evolution, but their role is underappreciated (5). The mutation spectrum is instantaneously and relatively easily altered by loss-of-function mutations in DNA repair genes, e.g., as observed in “mutators” that also have higher mutation rates (17). The elevated mutation rate increases the supply of beneficial mutations, facilitating rapid adaptation (18). However, spectrum shifts also occur without associated mutation rate changes, such as with the loss of some bacterial DNA repair enzymes (17), and under nutritional or anaerobic stress (19, 20). Diverse mutation spectra occur even across natural yeast strains and closely related species of *Chlamydomonas*, despite similar mutation rates (21, 22).

In fact, most species analysed so far have a skewed mutation spectrum (i.e., some mutational classes are over-represented compared to others), with substantial variation in the magnitude of bias (23, 24) (experimental data from microbes summarized in Table S1). This diversity implies major evolutionary shifts in mutation spectra. However, the frequency, underlying evolutionary processes, and consequences of spectrum shifts are unknown. For instance, do spectrum shifts evolve under selection? A spectrum shift may persist if the new bias favours mutational classes that are inherently advantageous (5, 25), allowing the evolved bias to hitchhike with beneficial mutations. It has also been speculated that stress-induced changes in mutation spectra may enhance sampling of new beneficial mutations by altering the distribution of fitness effects (DFE) (19, 26). However, prior studies have failed to find consistent support for general selective benefits of specific mutational classes. For instance, transition mutations (Ts) are not consistently more beneficial (or less deleterious) than transversions (Tv) (27, 28); and the fixation probability of Ts vs. Tv mutations in genomes is clade-specific (29). Hence, specific mutational classes do not seem to be universally beneficial, and the observed diversity in mutation bias remains puzzling.

We addressed these gaps using a combination of experiments, simulations, and phylogenetic analyses. To measure the immediate evolutionary impact of the mutation spectrum, we first estimated the genome-wide DFE of new mutations in *Escherichia coli*. We manipulated the mutation spectrum by deleting a DNA repair gene (creating a Δ*mutY* “mutator” strain) from our wild type strain (“WT”), altering both the mutation spectrum and rate (17). Allowing independent lineages of both strains to evolve under mutation accumulation (MA), we sequenced whole genomes of evolved isolates to identify strains carrying a single new mutation each. Thus, we minimized the impact of selection on the mutation spectrum and the DFE, while also decoupling the effects of the mutator’s spectrum from its high mutation rate. Measuring the effect of each single mutation on population growth rate, we determined the effects of mutation spectrum on the beneficial mutation supply, genetic load, and pleiotropic effects. Next, to test the generality and longer-term evolutionary consequences of shifts in mutation spectra, we performed both adaptive walk and full population simulations. Finally, we inferred evolutionary transitions in DNA repair enzymes across the bacterial phylogeny to estimate the frequency of changes in the direction of mutation bias. Our results show that shifts in mutation spectra can fuel adaptation via previously under-sampled evolutionary paths.

## RESULTS AND DISCUSSION

### The mutator DFE has more beneficial mutations that reduce the expected genetic load and alter pleiotropic effects

From MA experiments conducted in rich media (LB), we obtained 80 evolved WT (30) and 79 mutator strains, each carrying a single distinct mutation with respect to its ancestor (Supplementary Data). We measured the selective fitness effect (relative growth rate) of each mutation in the MA environment (LB) as well as 15 other environments with different carbon sources, including some that are used by *E. coli* in the mouse gut (31). With this large set of fitness effects, we constructed environment-specific empirical DFEs (WT data for some environments were previously reported (30)). Since bacterial MA experiments can cause over-sampling of beneficial mutations and under-sampling of deleterious mutations, we used a correction (32) that retains the measured selective effect (*s*) of each mutation, but estimates and corrects for selection bias by changing the frequency of mutations with a given selective effect (Fig. S1A–C, Supplementary Data). These corrected DFEs thus provide accurate estimates of the fraction of beneficial mutations (f_b_) and the average selective effect (*s*).

Correcting for selection bias during MA results in DFEs with coarsely binned fitness histograms, which we used to calculate the fraction of neutral (−0.05 < *s* < 0.05, conservatively accounting for a maximum of ~5% fitness measurement error), beneficial (*s* > 0.05), and deleterious (*s* < −0.05) mutations for each strain in all 16 environments. The DFEs and f_b_ values varied across environments (Fig. 1, Fig. S1), but the mutator often had a greater fraction of beneficial mutations and fewer deleterious mutations (Fig. 1, chi-square tests reported in Table S2; pooling data across all environments, average f_b_ = 36% vs. 28% for mutator vs. WT). Thus, compared to WT, the mutator had more beneficial-shifted DFEs. Mean fitness effects of mutations were also significantly more beneficial in the mutator (paired Welch’s two-sample test, p = 0.043; Fig. S2A), although medians were not significantly different (p = 0.35, Fig. S2B). Finally, mutations had larger absolute effects in WT than in the mutator (Wilcoxon’s rank-sum test, W = 662920, p = 1.4×10^-14^); hence, the mutator’s beneficial-shifted DFEs could not have arisen because of overall larger-effect mutations. These results indicate a broad selective advantage to the mutator, independent of its mutation rate. In addition, accounting for the skewed DFEs and order-of-magnitude higher mutation rate (μ_WT_ = 9.3×10^-11^ per bp per generation, μ_mutator_ = 89×10^-11^, Table S3), we estimated (following (33)) that on average the mutator should have an ~16-fold greater genome-wide supply of beneficial mutations (Table S4), and ~6-fold higher genetic load (number of deleterious mutations per genome) than the WT (Table S5). In contrast, if we ignored the mutator’s distinct DFE and accounted only for its mutation rate, both the supply of beneficial mutations and the deleterious genetic load would be nearly 10-fold greater than WT (Tables S4, S5; only the load is significantly affected by the corrected DFE). Thus, depending on their mutation rate and DFE, mutators may have a substantially higher supply of beneficial mutations and smaller genetic load than previously believed.

**Figure 1.**
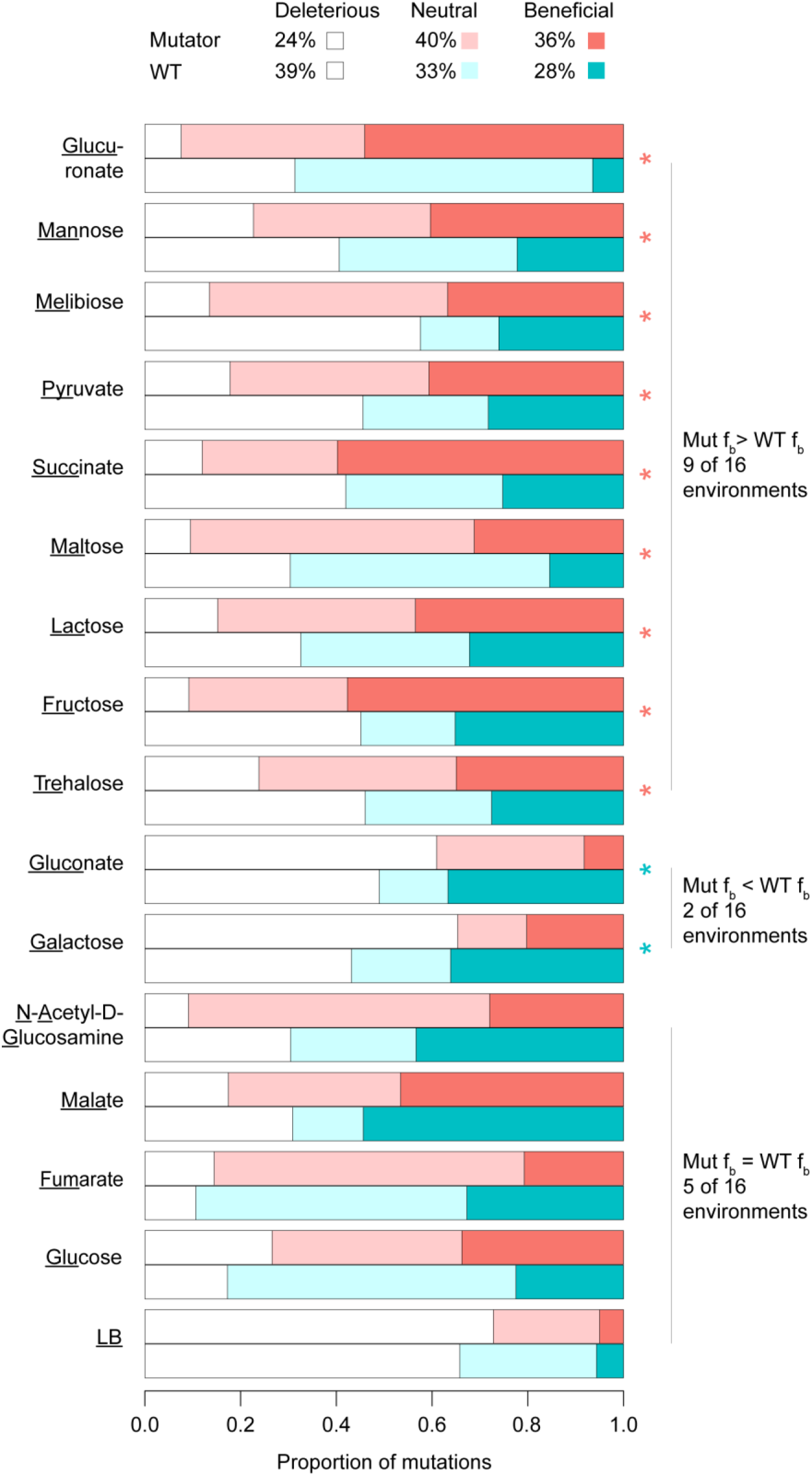
Mutator DFEs have a larger fraction of beneficial mutations than WT. Stacked bar plots show the observed proportion of beneficial (f_b_), deleterious and neutral mutations in WT and mutator (Mut). Average values for data pooled across environments are given in the key. Asterisks indicate significant differences between WT and mutator f_b_ in each environment, colored to show the strain with higher f_b_ (chi-square test output shown in Table S2).

The high f_b_ values observed in some environments (Fig. 1) contradict the general expectation that beneficial mutations should be rare (34). However, many other studies – including with microbes and plants – have also reported a large number of beneficial mutations (35–41). Such large f_b_ values have been partially attributed to an improved ability to measure small-effect beneficial mutations (38, 42, 43) that are more common than expected (44), especially at low effective population sizes (40). Most prior studies (34) provide approximate or indirect estimates of the DFE via random mutagenesis of a single gene, and focus only on the beneficial part of the DFE, or infer the DFE by fitting a model to time series data from selection or MA lines. Further, they do not estimate the number of underlying mutations to account for epistasis. Our direct, genome-wide estimates of the DFE of single, randomly-sampled new mutations support the idea (45, 46) that new beneficial mutations may be more common than generally believed.

Next, we estimated the pleiotropic effects of mutations (Fig. 2A), which are important because they can facilitate adaptation (via synergistic pleiotropic benefits) and shape fitness tradeoffs (via antagonistic pleiotropy) in new environments. The mutator had a distinct distribution of pleiotropic effects in all 16 environments (compare Fig. S2C–D, chi-square tests: p < 10^-5^ in each case, Table S6), characterized by a higher incidence of beneficial synergistic pleiotropy and lower deleterious synergistic pleiotropy. However, both strains had low antagonistic pleiotropy (Fig. 2B–C). Beneficial mutations in the mutator were also beneficial across many more environments (Kolmogorov-Smirnov test comparing the two distributions in Fig. 2F, D=0.41, p=3.1×10^-6^); whereas deleterious mutations were deleterious in fewer environments than WT (D=0.56, p=2.63×10^-11^; Fig. 2F, Fig. S3). Despite this qualitative association between mutational effects, overall, the magnitude of fitness effects was not correlated across environments (only 38 of 120 correlations were significant in WT and 22 of 120 significant in mutator, Spearman’s rank correlation, p < 0.05). Thus, new beneficial mutations in the mutator are more likely to facilitate adaptation across many environments, but are no more likely to generate tradeoffs. Interestingly, simulating an increase in the median fitness effect of the WT DFE without changing its shape (“WT-beneficial shift”) mimicked pleiotropic effects observed in the mutator in all but two environments (Fig. 2D, Fig. S2E, Table S6). Conversely, simply reducing the median fitness effect in the WT DFE (“WT-deleterious shift”) lowered beneficial synergistic pleiotropy in all 16 environments (Fig. 2E, Fig. S2F, Table S6). Ignoring the mutation spectrum can thus cause overestimation of a mutator’s genetic load, and underestimation of the supply and pleiotropic effects of beneficial mutations.

**Figure 2.**
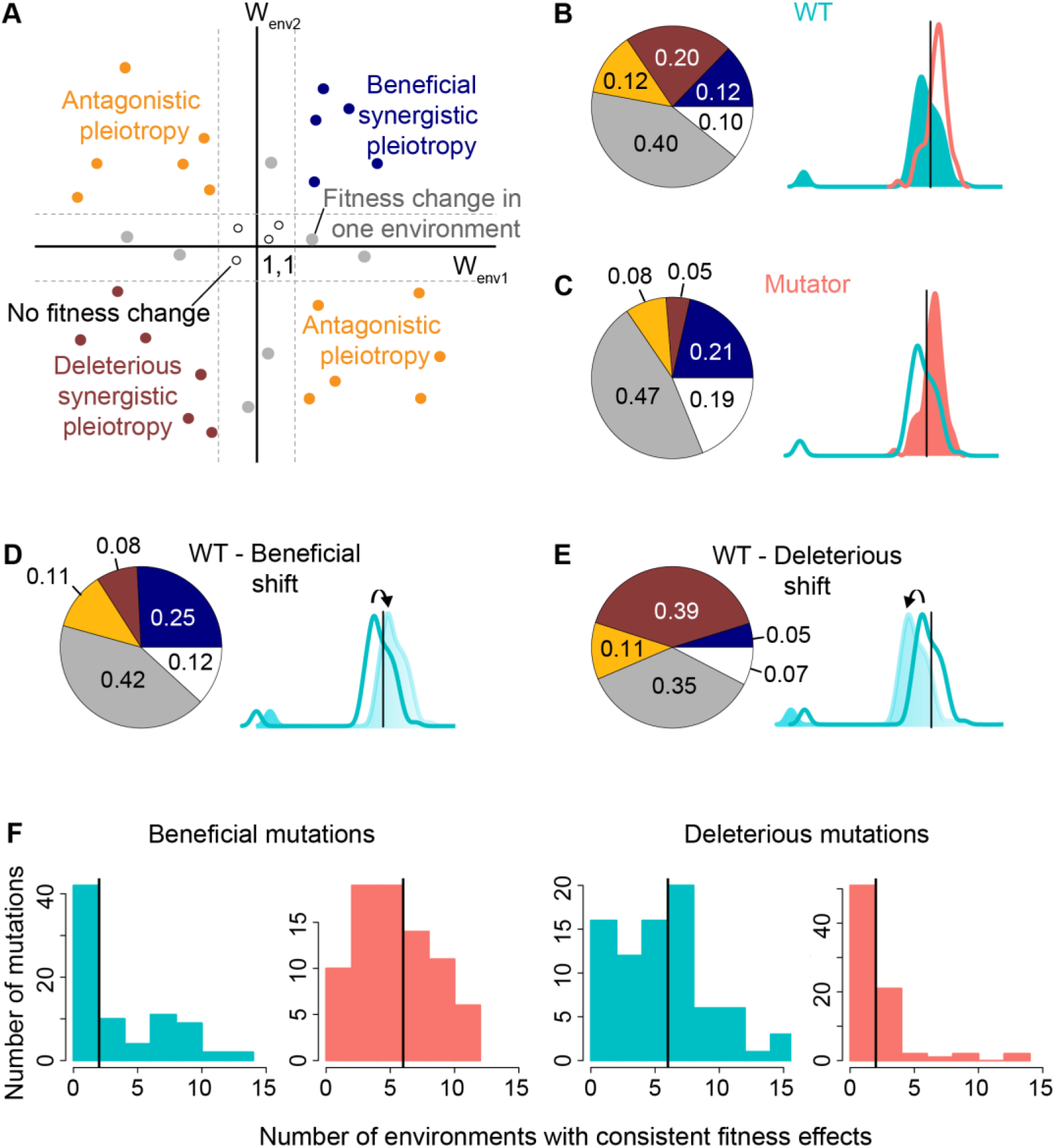
Beneficial-shifted DFE alters pleiotropic effects. (A) Schematic showing possible pleiotropic effects of mutations in two environments (W: relative fitness). (B–E) Median proportions of pleiotropic fitness effects (across 15 focal environments, excluding LB; also see Fig. S2C–F) of mutations in WT and mutator, and WT DFE artificially shifted towards either beneficial or deleterious mutations. Schematics indicate global DFEs (WT=cyan, mutator=pink, WT-shifted=light blue); black lines: relative fitness = 1. (F) Pleiotropic effects of mutations across multiple environments. Frequency distributions show the proportion of mutations with consistent fitness effects (beneficial or deleterious) in a given number of environments. Black lines indicate medians.

### The mutator’s beneficial-shifted DFE arises from its distinct mutation spectrum

We had hypothesized that the mutator’s distinct DFE could arise from its biased spectrum. However, other global effects of the initial *mutY* gene deletion could also lead to a DFE shift. For instance, if deleting *mutY* reduced the mutator’s fitness, a larger fraction of new mutations might be beneficial (47). However, the mutator ancestor had lower fitness than WT in only three environments (Fig. 3A, Table S7). Alternatively, epistatic interactions with *mutY* could increase the deleterious effect of new mutations in the WT. However, paired WT and mutator strains carrying the same mutation had similar fitness in all but one environment (Fig. 3B, Table S8). Thus, the observed shift in the mutator DFE cannot be explained by global effects of the *mutY* deletion. The difference across strains was also robust across mutational classes: mutations were more beneficial in the mutator even when comparing only base-pair substitutions, or coding, non-coding, synonymous, or non-synonymous mutations respectively (Fig. S4).

**Figure 3.**
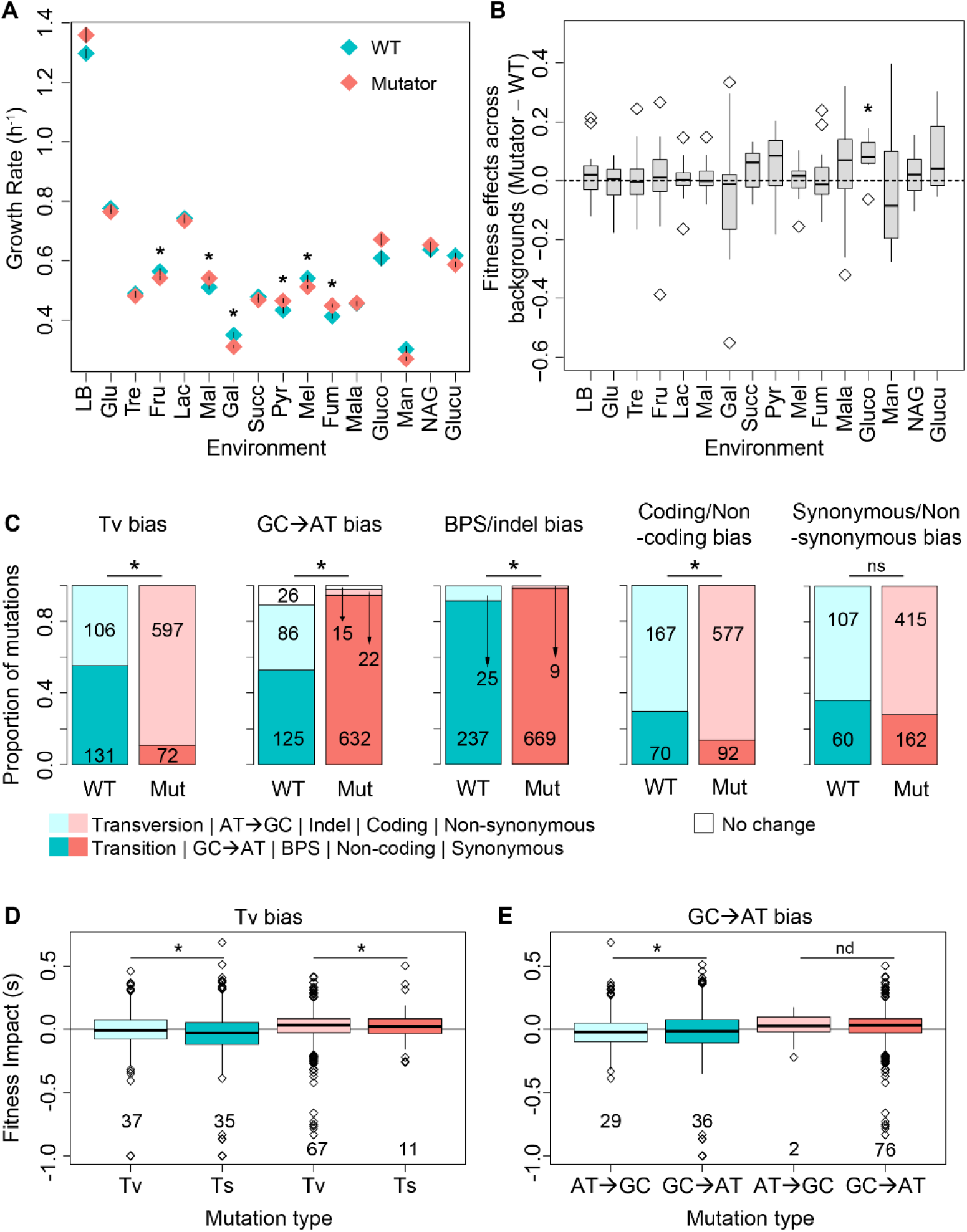
Differences in WT and mutator DFEs are best explained by differences in mutation spectra. (A) Growth rate of WT and mutator ancestors (n=12–24/strain/environment). Asterisks: significant differences between strains (Table S7). (B) Pairwise differences in the fitness effects of 19 mutations across genetic backgrounds (Mutator – WT); asterisks: significant difference from zero (black dotted line; Table S8). (C) Observed mutation biases in WT and mutator MA lines (including those with multiple mutations); the number of mutations of each type is noted. Asterisks: significant differences in proportions across strains (Table S3). (D–E) Fitness effects of different mutation classes, for MA-evolved isolates carrying single mutations. Boxplots (colored by strain and mutation class) show median, quartiles and range of relative fitness effects of mutations (open diamonds: outliers). Each boxplot was constructed with n x 16 data points (n = number of mutations, shown below each plot; pooled effects from 16 environments). Black horizontal lines: fitness impact = 0. Asterisks: significant differences across mutation classes (ANOVA output, Table S9; for imbalanced sample sizes, permutation tests as described for Fig. S4); nd: not determined.

Next, we asked whether the mutator is biased towards mutational classes that happen to be more beneficial (or less deleterious). We defined mutation bias as the proportion of mutations belonging to a specific mutational class (e.g., Tv bias = Number of Tv mutations / (Number of Tv + Number of Ts mutations)). Regardless of the type of bias, we could compare mutation biases across the new mutations sampled by WT and mutator, because barring the deletion of *mutY* both ancestral genomes were identical. The mutator strongly favours transversions, GC→AT mutations, base pair substitutions (BPS) and coding mutations (Fig. 3C, Table S3). Interestingly, in the WT strain, Tv and GC→AT mutations were more likely to be beneficial (29% for both) than Ts and AT→GC mutations (25% beneficial each). Overall, the median fitness effect of transversions and GC→AT mutations was less deleterious than transitions and AT→GC mutations respectively (Fig. 3D–E; fitness effects pooled across environments, Table S9. Note that excluding lethal mutations in WT – all of which were transversions – would enhance this effect. Importantly, Tv and GC→AT mutations had similar fitness effects in both strains (permutation tests: 61% and 85% of subsamples showed no difference between WT and mutator for Tv and GC→AT respectively; excluding lethal mutations: 73% and 87% of subsamples showed no difference between WT and mutator for Tv and GC→AT respectively). Other mutation biases did not show consistent differences (Fig. S4, Table S9). Additionally, other aspects of the mutations (e.g., genomic location or specific amino acid changes) also did not differ across strains (Fig. S5, S6; Tables S10, S11). Thus, as predicted, the mutation spectrum – specifically the strong Tv and GC→AT bias – appeared to drive the distinct DFE of the mutator, because the mutator samples these specific mutational classes more than the WT. However, it was unclear why Tv and GC→AT mutations should be more beneficial.

### Simulations demonstrate a general benefit of reversing mutation spectra

To test the generality of our empirical results and uncover underlying mechanisms, we simulated adaptive walks in an NK fitness landscape ((48), see Methods), modelling sequences of N nucleotides such that mutations could be classified as Ts vs. Tv or GC→AT vs. AT→GC. Starting from a randomly chosen ancestor sequence and mutation spectrum (Tv bias = fraction of transversions), we allowed populations to explore successive mutational steps at randomly chosen loci (bases). A mutation that confers a relative fitness increase of 1+s fixes (a step in the walk is taken) with probability 2s; that is, we assumed a strong-selection weak-mutation (SSWM) regime. At various points during the walk, we simulated mutations to generate a DFE, computed fitness effects given the underlying fitness landscape (affected by K other randomly chosen loci, to incorporate epistasis), and calculated the fraction of beneficial (f_b_) and deleterious (f_d_) mutations and their effect sizes. We initially set N=200 and K=1 (each locus epistatically coupled to 1/199=0.5% loci), and present the average outcomes for 500 adaptive walks on 500 randomly-generated fitness landscapes (Fig. S7A–B shows variation across walks).

As expected, mean population fitness increased during the adaptive walk, with a concomitant reduction in f_b_ and beneficial effect size, an increase in f_d_, and a relatively constant deleterious effect size (Fig. 4A). Setting ancestral Tv bias to 0.45 (mimicking our WT), we compared the DFE generated by the ancestor at various time points, to the DFE that would be created if the bias were changed at that time (mimicking our Δ*mutY* strain). The bias-shifted (“shifted”) strain thus has the same fitness and sequence as the ancestor and an identical mutation rate, differing only in spectrum. As the ancestor evolved (i.e., f_b_ decreased), a stronger bias shift (i.e., higher Tv) allowed the bias-shifted strain to sample proportionally more beneficial mutations than the ancestor (Fig. 4B). More generally, exploring well-adapted populations (f_b_ = 0.04) but varying both the ancestral and shifted mutation spectra, we find that shifted strains that reduce or reverse the ancestral bias (i.e., increase the sampling of mutations that were under-sampled by the ancestor, Fig. 4B) have the greatest advantage (Fig. S8). Our results hold across a different axis of the mutation spectrum (GC→AT bias, Fig. S9), and in more rugged fitness landscapes with higher epistasis (Fig. S10A–B). Importantly, adapting populations did not “exhaust” a small set of beneficial mutations; rather, they fixed distinct mutations as they explored a large number of diverse pathways that represent a subset of many possible fitness peaks (Fig. S10C). Our results also hold in full population simulations with multiple segregating mutations (relaxing the SSWM assumption), where deleterious mutations can fix (Fig. 4C–D, Fig. S11). In fact, when more mutations are accessible, we observe the same beneficial effects of bias shifts, but these effects emerge more quickly.

**Figure 4.**
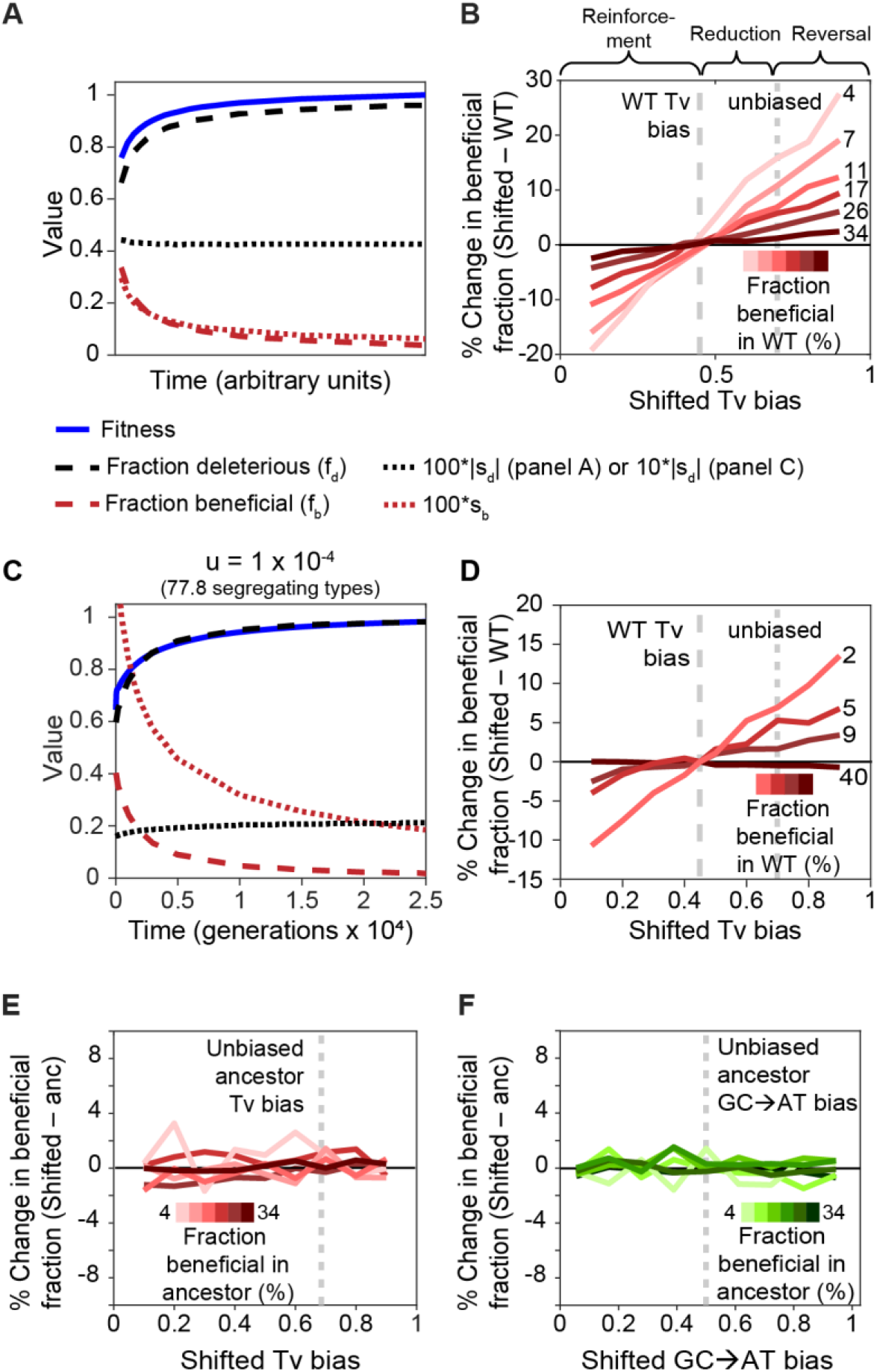
Simulations show a general benefit of reversing mutation spectra. (A) Change in mean population fitness, fraction of beneficial and deleterious mutations (f_b_ and f_d_), and mean magnitude of beneficial and deleterious effects (sb and sd) over the course of adaptive walks of a WT ancestor. (B) Impact of altering the WT mutation bias on f_b_ at different points along the adaptive walk (Tv bias = Tv/(Ts+Tv); grey dashed line: WT bias), as a function of the new mutation bias (shifted Tv bias). Reinforcements shift an existing bias further away from the unbiased state in the same direction. Reductions shift towards the unbiased state but not past it, and reversals shift the bias towards and past the unbiased state. Line color: WT f_b_ at a given time; steeper lines (lighter colors): later evolutionary time, better adapted WT. This figure shows simulation results for an ancestral Tv bias of 0.45; results for the full range of possible ancestral Tv biases (0.1 to 0.9) are shown in Fig. S8. (C–D) Results as in (A–B) but for simulated populations of 100,000 individuals evolving for up to 100,000 generations with a mutation rate of 10^-4^ per genome per generation; the mean number of segregating genotypes in each generation was 77.8. (E–F) Impact of introducing a mutation bias in an unbiased ancestor (“anc”), as a function of the new bias (shifted Tv or GC→AT bias). Panels show the results of simulations identical to those described in panel B, but starting with an unbiased ancestor.

Finally, these effects are also observed in full population simulations when the fitness landscape is defined using a codon-based rather than a nucleotide model (Fig. S12). Importantly, our simulation results imply that the benefit of a specific mutation class (e.g., Tv) is not universal, but is context-dependent: increased sampling of any mutational class that was previously poorly explored is beneficial.

Thus, our simulation results can be broadly generalized in the form of three key predictions: (1) reducing or reversing an ancestral mutation bias is always beneficial (2) the magnitude of the benefit increases with the degree to which the new bias opposes the ancestral bias, and (3) the magnitude of the benefit increases when the ancestor has evolved with a given bias for longer intervals of time (i.e., as the ancestor gets better-adapted). Our experimental data with the Δ*mutY* strain support the first prediction, since we found that overall, the mutator (with a reversal of the Tv bias, i.e., sampling many more transversions than the WT ancestor, Fig. 3D) had higher f_b_ values than WT. Interestingly, the WT GC→AT bias was reinforced in the mutator, rendering the results in Fig. 3E puzzling. However, ~88% of all GC→AT mutations in the mutator are also transversions. Hence, our mutator’s observed beneficial-shifted DFE is explained by its strong Tv bias, which opposes the existing WT bias towards Ts.

Although our data are generally explained by the simulation results, some puzzling observations remain. The simulations predict that the mutator should have higher f_b_ in all environments, but we do not observe this pattern as consistently as expected (Fig. 1). Further, in apparent contradiction to prediction 3 above, the difference in f_b_ between WT and mutator in lowest in LB (where ancestral growth rate is highest and presumably *E. coli* is best-adapted). However, note that our strains evolved for a very short time in LB under minimal selection, so LB is not an “ancestral” environment in an evolutionarily meaningful sense. Additionally, absolute growth rate is not a reliable indicator of the relative degree of or period of adaptation to an environment, because it is also strongly influenced by nutrient availability. Indeed, f_b_ was not correlated with ancestral growth rate across environments in either strain (Spearman’s rank correlation test, WT: rho = −0.217, Mutator: rho = −0.18; p > 0.05 in each case). Hence, to understand the environment-specific differences in DFEs, we need to explicitly test predictions 2 and 3 (described above) under conditions where the precise degree of bias change and the length of evolutionary time in an environment can be measured and/or manipulated.

Our simulation results demonstrate general conditions under which a change in mutational bias can be selectively advantageous. But how does mutation bias emerge in the first place? To address this, we note that an unbiased spectrum, by definition, samples all possible states in the state space with equal probability. The unbiased Tv fraction is thus 0.67 because 2/3^rd^ of all possible single nucleotide changes are transversions. For GC→AT bias, in the randomly generated genomes in our simulations, the unbiased spectrum is 0.5, while for genomes with extreme GC content, the unbiased state may change (e.g., in a GC-rich genome, the rate of GC→AT mutations would need to be higher to sample every possible alternative state with equal probability). Starting from an unbiased ancestor, we find that introducing mutation bias does not, on average, change the beneficial fraction of the DFE (Fig. 4E–F). Thus, if an ancestor could somehow achieve an unbiased state, there is no consistent selective pressure that would oppose the subsequent introduction of bias. In fact, in particular unbiased walks, introducing mutation bias can improve access to beneficial mutations, when by chance more of the remaining beneficial mutations are of a particular type (Fig. S7C); in these cases selection would drive the spectrum away from the unbiased state. We emphasize however that this unbiased state is distinct from a “null” expectation that may reflect various biochemical and enzymatic processes that influence mutation biases. The exact impact of these processes remains unclear and varies across organisms, such that a biologically meaningful and generalizable null expectation is not easily defined (49, 50). Nonetheless, after evolution under any existing bias, we predict that changes to the DFE that reduce or reverse the bias are favoured, allowing the population to explore mutational space that was previously under-sampled, and thus increasing the probability of finding new beneficial mutations. After a period of evolving with a new spectrum, a change in the fitness landscape (e.g., due to epistasis or environmental change) may again render a spectrum shift advantageous. We reiterate that larger magnitude shifts are more strongly favoured. Importantly, such large magnitude reversals of the existing bias that “overshoot” the unbiased state (i.e., Tv bias = 0.67 and GC→AT bias = 0.5) would allow even greater access to beneficial mutations than simply reducing the existing bias (Fig. 4B and 4D). Such large shifts can occur easily via a few mutations that lead to the gain or loss of DNA repair function (17). Together, these results suggest that evolutionary shifts in mutation spectra may be common, and should be enriched for bias reductions and reversals (i.e., bias shifts that improve the sampling of previously under-sampled mutational classes). We tested this prediction using a broad phylogenetic analysis of bacterial taxa.

### Evolutionary transitions in DNA repair enzymes are consistent with mutation spectrum reversals in most bacterial lineages

As noted earlier, altering DNA repair function is a simple and instantaneous mechanism to generate spectrum shifts. Hence, we asked whether long-term patterns of gain and loss of DNA repair genes in bacterial lineages support our prediction that a reduction or reversal in mutation bias is advantageous and should therefore occur more frequently than bias reinforcements. A direct test would require knowledge of the mutation bias of many extant taxa. Given the paucity of such data, we tested for qualitative patterns of evolutionary change. If successive evolutionary transitions in repair genes (i.e., gains or losses) typically lead to bias changes in opposite directions, this would be consistent with our prediction. For example, if an initial loss of a repair gene increases Tv mutations, we predicted that a subsequent evolutionary transition should lead to reduced Tv rather than a further increase in Tv (see Fig. 5A for possible phylogenetic outcomes and implications). We focused on 11 bacterial DNA repair enzymes whose deletion has known impacts on the *E. coli* mutation spectrum (Table S12). Assuming that the qualitative effects of deleting a specific enzyme are consistent across species (supported by available experimental data, Table S13), we inferred the direction of change in mutation bias following each gain or loss event in a lineage (e.g., the loss of *mutY* should increase Tv, and a subsequent loss of *mutT* will further increase Tv, so these successive events change bias in the same direction).

**Figure 5.**
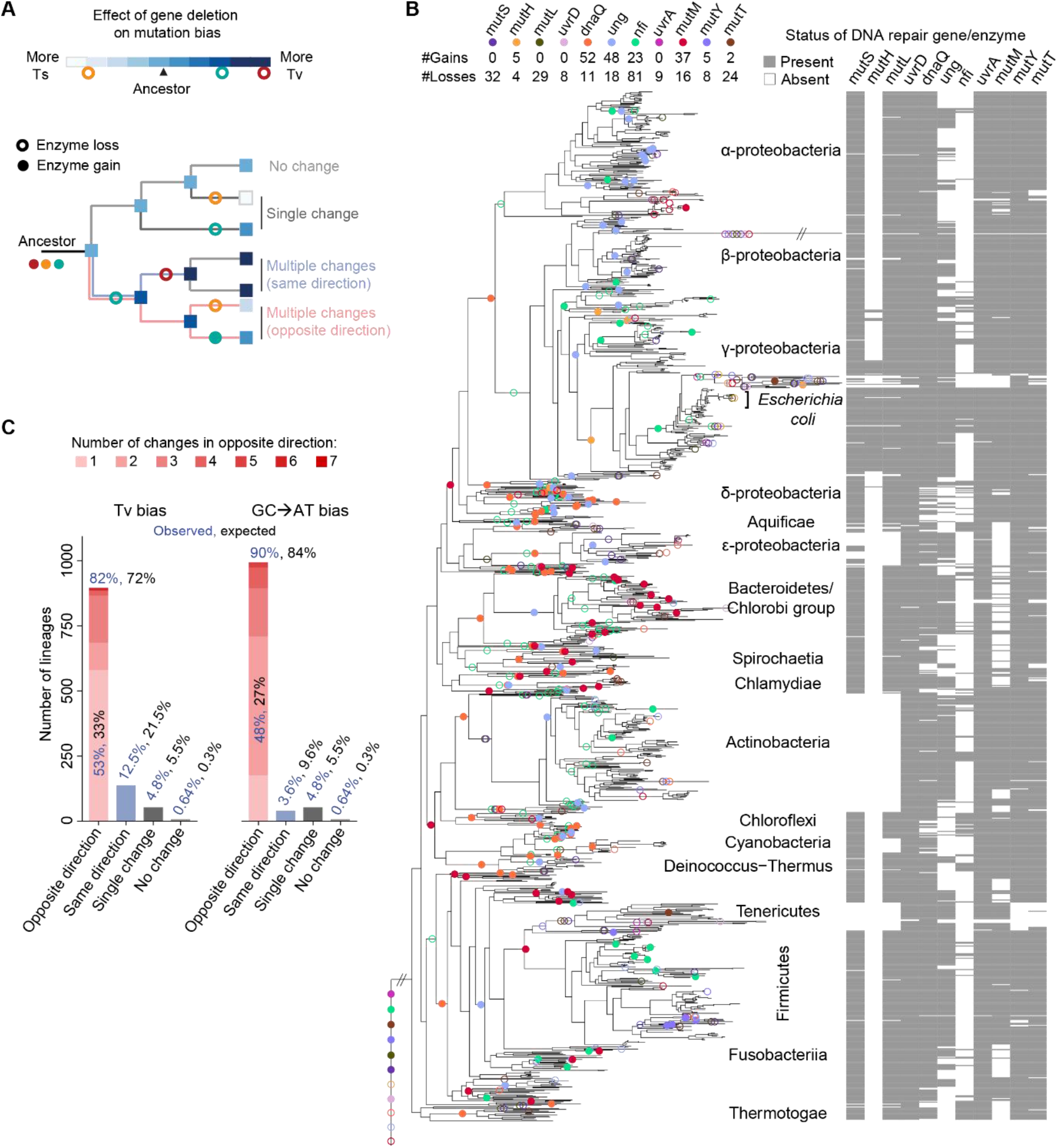
Phylogenetic analysis shows frequent evolutionary transitions in DNA repair genes and mutation bias. (A) Schematic showing the steps involved in the inference of changes in the direction of mutation bias in bacterial lineages. Different DNA repair enzymes are indicated by distinct colours; the qualitative effect of the loss of each gene on the mutation bias (based on experimental data from *E. coli*, Table S12) is shown in the color key. Separately, the gain and loss of each enzyme (indicated by open or closed circles) is inferred using ancestral reconstruction. Combining the two pieces of information, for each lineage we classify the number and type of changes as shown. Cases of multiple changes allow us to test our prediction that successive events should change mutation bias in opposite directions. (B) Repair gene presence/absence in extant taxa, used to map evolutionary transitions in DNA repair genes on the bacterial phylogeny (total number of events are shown in the key; see Supplementary methods for further details). (C) Observed number of lineages that show distinct types of evolutionary transitions in the direction of mutation bias (% values show fraction across all lineages; blue=observed; black=expected).

We first identified orthologs for each repair gene in 1093 extant bacterial genomes, and then used ancestral reconstruction to infer enzyme gains and losses on a phylogeny of these bacteria (following (51)) (Fig. 5B). We found frequent evolutionary transitions in most repair enzymes, including those with known effects on mutation bias (Fig. 5B) and 33 other genes with unknown effects on mutation bias (Fig. S13). For genes with known effects, we traced successive enzyme gains and losses for each lineage in the tree, and inferred the resulting fluctuations in the direction of bias change (Fig. 5A). Consistent with our prediction, in most lineages (>80%) successive enzyme gain or loss events would change mutation bias in opposite directions, and we observed only a few cases where successive events would lead to bias shifts in the same direction (Fig. 5C). Successive bias shifts in opposite directions were observed more frequently than expected by chance (one-sample t-tests, Tv bias: t = −72.38, p<2×10^-16^; GC→AT bias: t = −59.42, p<2×10^-16^) while bias shifts in the same direction were less frequent than expected (Tv bias: t = −72.38, p<2×10^-16^; GC→AT bias: t = −59.42, p<2×10^-16^) (Fig. S14B–C). Hence, our results cannot be attributed to peculiarities of the enzyme set, tree topology, or number of evolutionary transitions. The rarity of lineages showing either no change or a single change in mutation bias (~5% of total, Fig. 5C) further suggests that bias shifts occur very frequently, consistent with prior phylogenetic analyses of DNA repair genes in bacteria as well as fungi (52, 53, 54). Presently it is unclear whether these shifts are driven by selection, and whether such selection acts on mutation rate or bias (or both). Nonetheless, our analysis suggests that successive evolutionary transitions in DNA repair genes typically change the bias in opposite directions, consistent with our broader prediction that such shifts should be selectively advantageous.

We note here a few limitations of this analysis. First, reconstruction of ancestral character states in phylogenetic datasets does not account for uncertainty in the underlying phylogeny (55). Second, the estimated direction of bias change does not account for other repair enzymes whose effects on the mutation spectrum are unknown, or for variation in the impact of enzyme loss or gain. Finally, we also likely underestimate the actual number of bias shifts because we could not include cases where repair function may be altered without gene deletion (e.g., due to loss-of-function point mutations). Nonetheless, our results are consistent with our broad predictions, and indicate that the long-term causes and impacts of shifts in mutation bias deserve further investigation.

## CONCLUSIONS

We show that shifts in the strength and direction of bias offer access to previously under-sampled beneficial mutations. Thus, a species’ evolutionary history may be critical in explaining the evolution of genomic bias (e.g., in base composition), without invoking particular underlying mechanisms or selection favouring specific mutational classes. Previous work showed that the transversion-biased mutators Δ*mutT* and Δ*mutY* have distinct DFEs for antibiotic resistance compared to WT, but the effect was attributed to transversions being more beneficial for the specific antibiotic target (13). However, by recasting the puzzle of mutational biases as a problem of searching a vast and dynamic mutational space, our model allows broad generalization across mutation spectra. In addition, constraints imposed by biochemical or biophysical processes may also shape mutation biases, and could help explain the observed pervasive transition bias across taxa. However, from an evolutionary standpoint we suggest that the diversity of biases observed in nature is not surprising, since large magnitude shifts are selectively favoured, and may occur easily and frequently via changes in DNA repair function.

While it is well established that access to *<u>more</u>* mutations (through an increased mutation rate) is advantageous (18), here we demonstrate that accessing *<u>different</u>* mutations can likewise be beneficial. Our results have important implications for the evolution of mutator genotypes, in which shifts in mutation spectra and mutation rates are often (though not always) entangled. Mutator dynamics are governed by the genetic load and supply of beneficial mutations (18, 33, 56). In a new environment (away from fitness optima), these parameters are primarily driven by mutators’ high mutation rate (57) – particularly under strong selection (58) – allowing them to hitchhike with beneficial mutations (59). We show here that the genome-wide beneficial supply rate is strongly influenced by the mutation spectrum across environments, and this effect is strongest in well-adapted populations. Further, we predict that the advantage of a high mutation rate should be enhanced if accompanied by a reduction or reversal of the ancestral bias, but diminished otherwise (60). A recent simulation study also predicts that differences in the mutation spectrum of mutators can change the magnitude of deleterious mutations; such differences can facilitate the loss of mutators under selection (61). Thus, mutation rate and spectrum may jointly govern the observed rise, persistence, and fall of mutators under selection (13, 33, 62). Such effects may explain why mutators with relatively small increases in mutation rate are also abundant in natural bacterial populations (63), and deserve further attention (13, 61) with theoretical and experimental analyses to dissect the role of mutation rate vs. spectrum.

Together with studies showing that mutation biases are pervasive and influence the genetic basis of adaptation under diverse conditions (see Introduction), our results demonstrate that mutation spectra may be important drivers of adaptation and innovation under myriad scenarios. When the beneficial mutation supply is limited by the existing spectrum, an antiparallel (opposite direction on the same axis) or orthogonal jump (on a different axis of the spectrum) could enhance sampling of new beneficial mutations, facilitating rapid adaptation. Our phylogenetic analysis likely underestimated such evolutionary shifts in mutation spectra, which may occur even more frequently on shorter timescales via horizontal transfer and recombination (64), and drive polymorphism in spectra across natural bacterial isolates (63, 65). As a result, genomes may often be out of compositional equilibrium, with interesting implications for subsequent bias shifts and for evolutionary inferences that assume such equilibrium. Hence, further work is necessary to more robustly quantify the frequency of spectrum shifts. Finally, we predict multiple cascading effects of shifts in mutation spectra, including a reduction in the waiting time for beneficial mutations, decreased likelihood of mutational meltdown under genetic drift, and distinct genetic pathways of adaptation. We hope that future work will test these predictions.

## METHODS SUMMARY

Here, we provide a summary of the methods important to understand our study. Details are given in Supplementary Materials and Methods.

### Bacterial strains and mutation accumulation (MA) experiments

We obtained the wild-type (WT) *E. coli* K-12 MG1655 strain and mutator strains (Δ*mutY*, Δ*ung* and Δ*mutM* in *E. coli* K-12 BW25113) from the Coli Genetic Stock Centre (CGSC, Yale University). We moved the Δ*mutY*, Δ*ung* and Δ*mutM* loci into the WT strain using P1-phage transduction and confirmed the knockout using PCR and whole genome sequencing. To test whether genetic background alters the fitness effects of single mutations observed during mutation accumulation (MA), we deleted the *mutY* locus from 19 evolved WT MA lines using P1-phage transduction.

We founded WT, Δ*mutY*, Δ*ung* and Δ*mutM* MA lines from single colonies, and propagated them through daily single-colony bottlenecks on LB agar for 8250, 330, 4400 and 5775 generations respectively. The MA protocol minimizes the effect of selection, allowing us to sample a wide range of mutations largely independent of their fitness consequences.

### Whole-genome sequencing to identify single mutations and determine mutation spectra

We sequenced individual colonies from MA experiments to identify all clones carrying only a single mutation relative to the WT or Δ*mutY* ancestor, using the Illumina HiSeq 2500 platform. For each sample, we aligned quality-filtered reads to the NCBI reference *E. coli* K-12 MG1655 genome (RefSeq accession ID GCA_000005845.2) and generated a list of base-pair substitutions and short indels (<10bp). We calculated mutational biases as the fraction of each class of mutations observed in our evolved strains (Table S3).

### Constructing single-mutation DFEs

We measured growth rates of all evolved MA isolates with single mutations and their respective ancestors in 16 different environments, and estimated maximum growth rate as a proxy of fitness. For each isolate, we used the average growth rate of three technical replicates to calculate the fitness effect (*s*) as (growth rate of evolved isolate/growth rate of ancestor) – 1. We used *s* values of mutations to construct strain- and environment-specific distributions of fitness effects (DFE). Although selection is minimized in MA experiments, it is not entirely avoided. We corrected both WT and mutator DFEs for the effect of such selection as described recently (32), for each environment. Calculations are shown in the Supplementary Data file.

### Estimating genetic load, beneficial mutation supply, and pleiotropy

We used the mutation rate and f_b_ (estimated via the DFE) of WT and mutator in each environment to estimate the deleterious load and supply of beneficial mutations, as described earlier (33) (Tables S4 and S5). To predict the impact of the strain-specific DFEs on adaptation in new habitats, we estimated the incidence of antagonistic and synergistic pleiotropy among new mutations sampled by each strain. Results for WT *E. coli* in some environments were previously reported (30). We similarly estimated pleiotropy of the 79 single mutations that arose in the mutator (Δ*mutY*) background, and compared the incidence of pleiotropy across strains.

### Simulations to test the effect of mutation spectrum changes

We used an established evolutionary model to test whether and to what extent changes in mutation spectra are generally expected to affect the DFE. Following Stoltzfus (66), we simulated adaptive walks on the NK fitness landscape (48), modelling sequences composed of strings of four bases (A, C, G, T), allowing mutations to be classified as transitions or transversions, GC→AT or AT→GC, and so on. In an NK fitness landscape, each locus in the sequence is randomly assigned K neighbours from among all loci in the model. The fitness contribution of locus i (ω_i_) then depends on the state of that locus and the state of its neighbours. For computational efficiency we initially simulated adaptive walks (strong-selection weak-mutation regime). At various times during the adaptive walk, we generated a DFE by randomly sampling new mutations from the set of available single-step mutations and estimated key parameters such as f_b_.

Next, we relaxed a number of assumptions to test the generality of the simulation results. We investigated higher degrees of epistasis by varying both N and K. We also simulated full populations, in which large numbers of mutations simultaneously segregate, and deleterious mutations can fix. Finally, we implemented a codon-based model in which nucleotide sequences evolved as described above, but fitness was defined by an NK fitness landscape based on amino acid sequence.

### Inferring past evolutionary changes in mutation spectra

To determine the evolutionary history of each DNA repair gene, we detected orthologs and mapped predicted gain and loss events for each gene on a pruned version of a published phylogeny constructed using >400 proteins, as described previously (51). We determined the enzyme state at each node using a posterior probability threshold of 0.7. We used the predicted evolutionary transitions to count the total number of gains and losses of each enzyme in the phylogeny. For all lineages in the tree, we used gene loss data from *E. coli* to infer the direction of bias change (e.g. increase or decrease in Tv mutations) after each gain or loss event, and used these to calculate the observed number and direction of changes in bias in each lineage. For lineages with multiple evolutionary transitions, we asked whether successive gain or loss events changed a given mutation bias in the same or opposite direction, and counted the number of occurrences of each type of event (Fig. 5A). To derive the number of changes in bias expected by chance alone, we ran 10,000 phylogenetic stochastic forward simulations on the transition rate matrices (describing inferred rates of enzyme state transitions) and used the number and direction of mutation bias changes in each simulation to calculate the average proportion of lineages showing a given number and type of change (Fig. S14A).

## Acknowledgements

We thank Shyamsunder Buddh, Brian Charlesworth, Deborah Charlesworth, Shachi Gosavi, Joachim Krug, Krushnamegh Kunte, Saurabh Mahajan, Christopher Marx, and Mukund Thattai for discussion; Joshua Miranda for assistance with MA experiments; and the NGS facility at the Bangalore Life Sciences Cluster.

## Author contributions

MS designed and conducted experiments, analysed data, and drafted the manuscript. GDD designed and conducted phylogenetic analyses. BAB conducted experiments and analysed data. LMW obtained funding, designed and conducted simulation analyses, and drafted the manuscript. DA conceived and designed experiments and phylogenetic analyses, analysed data, obtained funding, and wrote the manuscript.

## Data and materials availability

All data used for experimental and phylogenetic analysis are available as Supplementary Information files. Code for simulations and phylogenetic analysis is available on Github (https://github.com/lmwahl/MutationSpectrum and https://github.com/gauravdiwan89/dfe_ms_phylo_analysis).

