## Supplementary Information for "Shifts in mutation spectra enhance access to beneficial mutations"

### SUPPLEMENTARY MATERIALS AND METHODS

#### Bacterial strains

We obtained the wild-type (WT) strain of *E. coli* K-12 MG1655 from the Coli Genetic Stock Centre (CGSC, Yale University), streaked it on LB (Luria Bertani) agar, and chose one colony at random as the WT ancestor for subsequent experiments. We similarly obtained the mutator strain of *E. coli* (Δ*mutY*) from the Keio collection (BW25113 strain background; (1)) of gene knockouts from the same stock centre. These gene knockouts were made by replacing open reading frames with a Kanamycin resistance cassette, such that removing the cassette generates an in-frame deletion of the gene. The design of gene deletion primers ensured that downstream genes were not disrupted due to polar effects (1). We moved the knockout locus from the BW25113 background into the MG1655 (WT) background using P1-phage transduction (2). We then removed the kanamycin resistance marker by transforming kanamycin-resistant transductants with pCP20, a plasmid carrying the flippase recombination gene and $Amp^R$ resistance marker. We grew ampicillin resistant transformants at 42 ºC in LB broth overnight to cure pCP20. We streaked out 10 µL of these cultures on LB plates. After 24 hours, we replica-plated several colonies on both kanamycin-LB agar plates and ampicillin-LB agar plates, to screen for the loss of both kanamycin and ampicillin resistance. We PCR-sequenced the knockout locus to confirm removal of the kanamycin cassette. We used the same protocol to generate Δ*ung* and Δ*mutM* strains to determine their mutation spectra.

To test whether genetic background alters the fitness effects of single mutations observed during mutation accumulation (MA), we deleted the *mutY* locus from 19 evolved WT MA lines (see below). We raised P1 phage on the Δ*mutY*::kanR strain in the MG1655 genetic background and infected evolved WT MA lines to obtain unmarked Δ*mutY* strains carrying the evolved WT mutation, as above. We confirmed the *mutY* deletion by sequencing the *mutY* locus. Thus, for each of the 19 mutations, we obtained paired strains with either a WT or Δ*mutY* genetic background.

#### Experimental evolution under mutation accumulation (MA)

We founded 38 WT MA lines from 38 different colonies (two lines per Petri plate) incubated at 37ºC, as described earlier (3). For each line, every 24 hours we streaked out a random colony (closest to a pre-marked spot) on a fresh LB agar plate. Every 4-5 days, we inoculated a part of the transferred colony in LB broth at 37ºC for 2-3 hours and froze 1 mL of the growing culture with 8% DMSO at -80 ºC. For the current study, we used stocks frozen on days 39 (~1072 generations), 104 (~2860 generations), 140 (~3850 generations), 200 (~5500 generations), 250 (~6875 generations) and 300 (~8250 generations). For the mutator, we similarly founded 300 MA lines, and evolved them for 12 days (~330 generations). This time was expected to be sufficient for the accumulation of one mutation per line on average, based on previous estimates of mutation rate (4). We made stocks of all lines as described for WT. To determine the impact of two previously untested DNA repair enzymes (*ung* and *mutM*) on the mutation spectrum, we evolved 20 independent MA lines of each deletion strain for 160 days (~4400 generations) and 210 days (~5775 generations) respectively.

#### Whole-genome sequencing to identify single mutations and determine mutation spectra

We sequenced individual colonies from MA experiments to identify all clones carrying only a single mutation relative to the WT or *ΔmutY* ancestor. For WT, we sequenced MA lines at days 39, 104, 140, 200, 250, 300 as described earlier (3), to obtain 80 single mutational steps that we used for the analyses in the present study. We sampled lines at different time points since each line acquired its first mutation at different time points. For *ΔmutY*, we sequenced whole genomes of the 300 colonies from the MA experiment frozen on day 12, and the Δ*mutY* MA ancestor. To do this, we inoculated 2 µL of the frozen stock of each evolved MA line (or the ancestor) in 2 mL LB, and allowed the cells to grow overnight at 37ºC with shaking at 200 rpm. We extracted genomic DNA (GenElute Bacterial Genomic DNA kit, Sigma-Aldrich), quantified it (Qubit HS dsDNA assay, Invitrogen), and prepared paired-end libraries from each line using the Illumina Nextera XT DNA library preparation kit per the manufacturer's instructions. We sequenced the Δ*mutY* MA ancestor on the Illumina MiSeq platform using the 2x250bp paired-end reaction chemistry, and obtained a total of 1.2M reads with quality > Q30, corresponding to an average per base coverage of ~120x. We sequenced libraries from the evolved MA lines on the Illumina HiSeq 2500 platform using the 2x100bp paired-end reaction chemistry. We obtained sufficient reads for 299 out of the 300 samples (median 3.3 million reads per sample; range: 1.15 – 7.13 million reads per sample), which we retained for further analysis. We discarded reads with quality < Q30, obtaining a median per base coverage of ~92x (range: 32x – 198x). For each sample, we aligned quality-filtered reads to the NCBI reference *E. coli* K-12 MG1655 genome (RefSeq accession ID GCA_000005845.2) using the Burrows-Wheeler short-read alignment tool (65). We generated pileup files using SAMtools (5) and used VARSCAN to extract a list of base-pair substitutions and short indels (<10bp) (6). For further analysis, we only retained mutations with >80% frequency that were represented by at least 5 reads on both strands.

After removing ancestral mutations from the evolved lines, we identified 79 *ΔmutY* MA-evolved lines that each had a single new mutation compared to the ancestor. We performed all further experiments and analyses with our 80 WT and 79 *ΔmutY* isolates (see Supplementary data). To calculate mutation rates and estimate mutation spectra, we used mutations called from all sequenced isolates at day 300 for WT and day 12 for Δ*mutY*, irrespective of whether they had single or multiple mutations (total 237 mutations in WT and 669 in mutator). Similarly, we determined the mutation spectra of Δ*ung* and Δ*mutM* strains by sequencing 20 isolates per strain (total 157 and 138 mutations respectively). We calculated mutational biases from the fraction of different types of mutations observed in our evolved strains (Table S3). We calculated the WT Tv bias as the number of transversions/(number of transitions + number of transversions) = 0.447. Therefore, we used WT Tv bias = 0.45 for our adaptive walk simulations (Fig. 4). Similarly, we calculated the WT GC→AT bias as the number of GC→AT mutations/(number of AT→GC mutations + number of GC→AT mutations) = 0.59 (ignoring 26 AT→TA and GC→CG mutations that did not contribute to the bias; Table S3). For convenience, in simulations we used the fraction of GC→AT mutations = 0.55 and the fraction of AT→GC mutations = 0.35, thus obtaining WT GC→AT bias = 0.61.

**Measuring growth rate as a fitness proxy**

We measured growth rates of evolved isolates with single mutations and their respective ancestors in liquid culture media: LB broth (Miller, Difco), or M9 minimal salts (Difco) + 5 mM carbon source (glucose, trehalose, fructose, lactose, maltose, galactose, succinate, pyruvate, melibiose, fumarate, malate gluconate, mannose, N-acetyl-D-glucosamine (NAG), or glucuronate; Sigma-Aldrich). We inoculated each isolate from its freezer stock into M9 minimal salts medium with 0.4% glucose, and allowed it to grow at 37 ºC with shaking at 200 rpm for 16 hrs. We inoculated 6 µL of this culture into 594 µL growth media in 48-well plates (Costar) and incubated in a shaking tower (Liconic) at 37 ºC. Plates were read by an automated growth measurement system (Tecan, Austria) every 35-45 minutes for 18 hrs. We measured the growth rate of three technical replicates per isolate per growth medium; meaningful biological replicates could not be obtained since we had a single colony at the end of each MA line. The technical replicates were inoculated in the same microplate and measured on the same day. In each 48-well plate, we included the WT ancestor to estimate variation in growth rates across plates. We estimated maximum growth rate, obtained from a linear fit to log (optical density) vs. time data, using the Curve Fitter software (7). Measurements across different days were highly repeatable (Fig. S1 in (3)) and measurements of the Δ*mutY* and WT ancestors in 12 environments were strongly correlated across two different years (in both cases, $R^2 > 0.99$, $p < 10^{-11}$).

For each evolved isolate, we used the average growth rate of three technical replicates to calculate relative growth rate as: Growth rate of evolved strain/Growth rate of ancestral strain. For WT MA lines, we used the WT ancestor; and for mutator MA lines, we used the mutator ancestor. The fitness effect of each mutation (*s*) is then calculated as relative growth rate –1, which is equivalent to (Growth $\text{Rate}_{evo}$ – Growth $\text{Rate}_{anc}$)/Growth $\text{Rate}_{anc}$. Usually, $s < 0$ indicates a deleterious mutation, while $s > 0$ indicates a beneficial mutation. However, the estimated error in measurement of growth rates across technical replicates (run on the same day) was at most 5%. Hence, we conservatively considered mutants with $s > -0.05$ and $s < 0.05$ as showing no change in fitness, or neutral. We also tried using a different method to calculate fitness effects (selection coefficients) of mutations, following (8): *s* = ((Growth $\text{Rate}_{evo}$ – Growth $\text{Rate}_{anc}$)/Growth $\text{Rate}_{anc}$)*ln2. We used *s* values of mutations to construct strain- and environment-specific distributions of fitness effects (DFE).

**Correcting DFEs to account for selection bias during mutation accumulation**

We corrected for selection bias during mutation accumulation as described recently (9). First, we corrected for selection bias in the MA environment (i.e., LB) for both WT and mutator. We binned observed fitness values in LB into 17 bins with a width of 0.05, to correspond with the maximum measurement error. Using the mean *s* value for each bin, we calculated the selection bias *b(s)* according to equation 2 (9). We used the $\tau$ value of 27 divisions per cycle of colony growth (i.e., per transfer in the MA experiment) as previously reported (3). We re-weighted each fitness bin by multiplying the bin frequency by $w = 1/b(s)$ to obtain the corrected frequencies in each bin. Finally, we normalized the corrected re-weighted frequencies as described in equation 3 (9).

Next, we corrected the DFEs in “new” environments (i.e., the 15 M9-minimal medium environments) according to equation 4 in (9). For each mutation, we used its fitness in LB to determine its weight $w_m = 1/b(s)$. Next, we binned fitness effects in the “new” environment into 17 bins, as done above for LB. In each bin, we calculated the sum of the $w_m$ values for all mutations whose fitness effects fell into that bin. Finally, taking these summed $w_m$ values as bin weights, we calculated bin frequencies, and re-normalized bin heights to sum to unity, as described above for LB. We performed these bias corrections for DFEs of both WT and mutator in each of the 15 new environments. The calculations are shown in the Supplementary Data file.

**Estimating expected genetic load and supply of beneficial mutations**

We calculated the deleterious load and supply of beneficial mutations for each strain as described earlier (10). Briefly, using the fraction of beneficial and deleterious mutations from the bias-corrected DFEs of WT and mutator, we calculated genetic load due to deleterious mutations as:

$$L_d = 1 - \exp(-U) \cong f_d \, N \, \mu$$

where $L_d$ = deleterious genetic load; U = deleterious mutation rate per genome; $f_d$ = fraction of deleterious mutations; N = number of sites (here, *E. coli* genome size, $4.64 \times 10^6$ bp); $\mu$ = average genome-wide mutation rate of each strain ($bp^{-1}$ $generation^{-1}$; Table S3). Similarly, we calculated the supply of beneficial mutations as:

$$S_b = f_b \, N \, \mu$$

where $S_b$ = beneficial supply and $f_b$ = fraction of beneficial mutations. Using the $L_d$ and $S_b$ values for each strain, we calculated the fold change in beneficial supply and genetic load of the mutator relative to WT. Next, we calculated the mutator $L_d$ and $S_b$ assuming that the WT and mutator DFEs were identical, and allowing for variation only in $\mu$, as done in previous studies:

$$S_{b(WT\ DFE)} = f_{b(WT)} \, N \, \mu_{\Delta mutY}$$

$$L_{d(WT\ DFE)} = f_{d(WT)} \, N \, \mu_{\Delta mutY}$$

To measure the impact of accounting for the mutator’s distinct DFE, we compared the fold change in the mutator’s $S_b$ and $L_d$ values compared to WT, with or without assuming distinct DFEs. All calculated values are given in Tables S4 and S5.

**Measuring the incidence of pleiotropy among new single mutations**

We estimated the incidence of pleiotropy among new single mutations in the mutator (Δ*mutY*) background, as reported earlier for single mutations in WT *E. coli* (3). Briefly, for each possible pair of environments (total 210 pairs, given 15 minimal medium environments), we calculated the proportion of new mutations that increased fitness in one environment while decreasing fitness in another (antagonistic pleiotropy), simultaneously increased or decreased fitness in both environments (synergistic pleiotropic increases or decreases), changed fitness only in one environment, or did not change fitness in either environment. For each focal environment, we then calculated the median proportion of each type of pleiotropic effect (considering all pairs that included that environment, excluding LB). Finally, for ease of visualization in Fig. 2, we calculated the median proportion of each type of pleiotropic effect across all focal environments, for both WT and mutator.

Next, we tested whether the differences in proportions of pleiotropy between WT and mutator could be recapitulated solely by shifting the WT DFE towards more beneficial mutations (“WT-beneficial shift”). We calculated the difference in median fitness effects of mutator and WT in each environment, and added this value to each of the WT fitness effects in that environment. The resulting DFE had a median equivalent to the mutator DFE, but retained the shape of the WT DFE. Similarly, we generated a WT DFE shifted towards more deleterious mutations (“WT-deleterious shift”) by subtracting the same value from the relative fitness values of WT mutations. We used these “shifted” DFEs to calculate the proportion of fitness effects showing each type of (pairwise) pleiotropy in the same way as described above for the WT and mutator. Finally, to measure pleiotropic effects across multiple environments, we counted the number of environments in which each mutation had a consistent effect (either beneficial or deleterious); and tested whether the proportion of mutations with consistent effects differed significantly across WT and mutator.

**Estimating the effect of mutation spectrum changes via adaptive walk and full population simulations**

We used an established evolutionary model to test whether and to what extent changes in mutation spectra are generally expected to affect the DFE. Following Stoltzfus (11), we simulated adaptive walks on the NK fitness landscape (12), modelling sequences composed of strings of four bases (A, C, G, T), allowing mutations to be classified as transitions or transversions, GC→AT or AT→GC, and so on. In an NK fitness landscape, each locus in the sequence is randomly assigned K neighbours from among all loci in the model. The fitness contribution of locus i ($\omega_i$) then depends on the state of that locus and the state of its neighbours. To create the fitness landscape when K=1, for each locus, 4^2 = 16 possible values of $\omega_i$ are drawn from a uniform distribution on (0,1); these values define $\omega_i$ if the state of the locus and its neighbour are AA, AC, AG... through TT. The fitness of the sequence (W) is then given by the sum of contributions from each locus, $W = \sum_{i=1}^{N} \omega_i$. We used neighbourhood sizes of K=1 to 8, corresponding to relatively smooth and relatively rugged fitness landscapes. When N=200, each landscape contained $4^{200} = 2.58 \times 10^{120}$ unique sequences and fitness values. Increasing K increases the number of local fitness peaks; for example, when N=20 and K=8, the landscape has approximately 7000 distinct local peaks (13). For computational efficiency we initially simulated adaptive walks (strong-selection weak-mutation regime).

When simulating Tv bias, starting at a randomly chosen resident (ancestor) sequence, we chose a single locus at random for mutation. The mutation is a transition with probability Ts and a transversion with probability Tv = 1 – Ts. Note that within transversions, each of the two possible outcomes (e.g., A→C or A→T) is chosen uniformly at random. The selective effect of the mutation was computed as s = $W_{mu}/W - 1$, where $W_{mu}$ is the fitness of the mutant sequence, and W is the fitness of the current population sequence. The mutant sequence replaces the population sequence (a “step” in the walk occurs) with probability 2s if s > 0. The mutation spectrum (value of Ts) does not change during the adaptive walk.

We pause the adaptive walk at various time points to create a DFE. To create the WT DFE, we generate 500 single-step mutants from the WT sequence, using the WT bias to generate the mutations. We then temporarily shift the bias to a new value, and create 500 random single-step mutants from the WT sequence using this shifted probability to generate the mutations. This process creates a single shifted-bias DFE; we then repeat this process for values of the shifted bias between 0 and 1.

We extended these simulations in a number of ways. We implemented an alternative mutation scheme to test if the results varied for GC→AT biased mutation spectra. In these simulations, mutations were constrained such that 10% were AT→TA or GC→CG (mimicking our WT), and of the remaining mutations, fraction *x* were AT→GC, while (0.9-*x*) were GC→AT. For this mutation scheme, GC→AT bias was calculated as the fraction of GC→AT mutations among all AT→GC or GC→AT mutations (Fig. S9). To investigate higher degrees of epistasis (more rugged landscapes), we varied both N and K, simulating adaptive walks and DFEs for N=8, 20 and 200, with K = 1-8 (Fig. S10).

Next, we conducted full population simulations instead of adaptive walks, allowing populations with carrying capacities of 100,000 individuals evolved for 100,000 generations. Each individual in the population reproduces with Poisson-distributed offspring, proportional to fitness, normalized by the population mean fitness. The carrying capacity is implemented using the Ricker model, in particular the mean number of offspring for every individual in a given generation is multiplied by exp(1-Nc/κ) where Nc is the current population size and κ is the carrying capacity. Each population is initialized with 50 random genotypes of 2000 individuals each. Fitness is determined as described previously using the NK model, with N=100 and K=5. The population evolves with a fixed Tv bias (0.45). At various time points during adaptation, we pause the adaptation process and find the most common genotype in the population. We then create 500 random single-step mutants of the most common genotype, again using the fixed Tv bias to create these mutations. We compute the fitnesses of these 500 mutants to determine the DFE in a given generation.

We used population simulations to examine the impact of relaxing the SSWM regime, by increasing the mutation rate and allowing large numbers of mutations to segregate simultaneously. In these simulations, deleterious mutations could also fix (Fig. S11). We also used the full population simulations to examine the impact of using a codon-specific model in which nucleotide sequences evolved as described above, but fitness was defined by an NK fitness landscape based on amino acid sequence (Fig. S12).

**Inferring past evolutionary changes in mutation spectra**

To determine the evolutionary history of focal DNA repair genes (11 with known effects on mutation bias and 33 with unknown effects), we detected orthologs and mapped predicted gain and loss events for each gene across the bacterial phylogeny, as described previously (14). Briefly, we pruned a previously described whole genome phylogeny based on >400 proteins (15) to generate a tree that included 1093 fully sequenced bacteria (14). Next, we downloaded the reviewed sequences for each DNA repair enzyme gene from UniProt, built a Hidden Markov Model (16), and searched for significant orthologues in the genomes of the 1093 bacteria. Finally, we noted the presence/absence of each gene in each extant genome and carried out ancestral reconstruction of gene state (presence/absence), to infer the enzyme state at all nodes in the phylogeny. We used stochastic character mapping (17) in the R package 'phytools' (18) with the following parameters: transition rate matrix determination – Bayesian MCMC, prior distribution on the root node of the tree – estimated, number of simulations – 500. These parameters allow the function to first estimate the probability of the ancestral state at the root of the tree, and then determine the transition rate matrix that best explains the gene presence/absence data at the tips. A stochastic map is constructed using this transition rate matrix, which gives the probability of each character state at every node in the phylogeny. The function was simulated 500 times, each generating a different transition rate matrix and probabilities at the nodes. To score evolutionary transitions in character states, we first assigned each node with a state based on an average posterior probability of >=0.7 for either presence or absence of the gene (i.e., nodes with a probability of <0.7 were assigned the state of the parental node; the root node was assigned the state with the higher probability), and then determined the nodes where the states switched. We used the predicted evolutionary transitions to count the total number of gains and losses of each enzyme in the phylogeny.

For each lineage in the tree, we determined the evolutionary sequence of gene gain and loss, from the root up to the extant taxon. Given this sequence of events, we categorized lineages as showing "no change", "single change", or "multiple changes" in the mutation bias (see schematic in Fig. 5A). To do so, we used empirical data on the change in mutation bias on deletion of specific DNA repair genes in *E. coli* (Table S12), classifying the impact of each gene loss as changing the bias in one direction (i.e., loss of any enzyme that caused an increase in Tv or an increase in GC→AT) or changing the bias in the other direction (i.e., loss of a gene that increased Ts or AT→GC). When available, we used experimental data on double or triple knockouts of DNA repair mutants to infer the direction of bias change when multiple enzymes were lost (Table S12). Otherwise, we assumed additive effects of the lost enzymes; e.g., if both *mutT* and *mutY* were lost, we would expect a larger increase in Tv mutations compared to a single deletion alone. Note that because the exact magnitude of bias shifts cannot be inferred without empirical measurements, we restricted our analysis to qualitative changes in the direction of bias.

To test whether the patterns of changes in the direction of bias were consistent with predictions from our adaptive walk simulations, we focused on lineages showing multiple changes (i.e., at least two gain/loss events predicted to impact mutation bias). For these cases, we asked whether the impact of successive gene gain/loss events would result in changes in bias in the same or the opposite direction. For instance, across nodes in a lineage, if one bias change increased Ts and the next change reduced Ts, this was counted as a change in the opposite direction. Thus, the total number of events of each type depends only on the direction of the bias change associated with each gene loss (e.g., more Tv or more Ts). Finally, we calculated the observed proportion of all lineages that showed successive changes in bias in the same vs. opposite direction.

To derive the expected frequency of each category of change in the direction of mutation bias based on chance alone, we extracted the 500 estimated transition rate matrices (describing inferred rates of enzyme state transitions) from the stochastic character mapping described above. We used the function 'sim.history' in the R package phytools (18) to run 20 forward simulations for each matrix, yielding a total of 10,000 evolutionary simulations where enzyme state transitions occurred with similar rates as observed, but at stochastically determined nodes in the phylogeny. As described above, we calculated the average proportion of lineages showing a given number of successive bias shifts in opposite directions (or other types of change in the direction of mutation bias) (Fig. S14A).

To validate our method of inferring the direction of mutation bias shifts given each species' (or hypothetical ancestral node's) complement of DNA repair enzymes, we asked if there is any relationship between the inferred *magnitude* of mutation biases for extant taxa and observed mutation biases from mutation accumulation studies (available data shown in Table S1; where multiple experimental measures exist, we used means). Given the paucity of experimental data, this analysis is likely under-powered; but the results suggest that our phylogenetic analyses are able to broadly capture the direction of actual biases. The observed and inferred Tv biases were significantly positively correlated (linear regression, $R^2 = 0.6187$, $p = 0.012$), and for GC→AT biases we found a marginally significant correlation (linear regression, $R^2 = 0.41$, $p = 0.049$ after removing an obvious outlier, *Deinococcus radiodurans*).

### SUPPLEMENTARY FIGURES

**Figure S1. Raw and bias-corrected DFEs of WT and mutator strain.** (A) Raw (gray) and corrected DFEs (cyan) of MA-accumulated mutations in WT, in each of 16 environments. (B) Raw (gray) and corrected DFEs (pink) of MA-accumulated mutations in the mutator background. (C) Corrected DFEs of WT and mutator in each environment. In panels A and B, in most cases, the raw and corrected DFEs overlap almost completely.

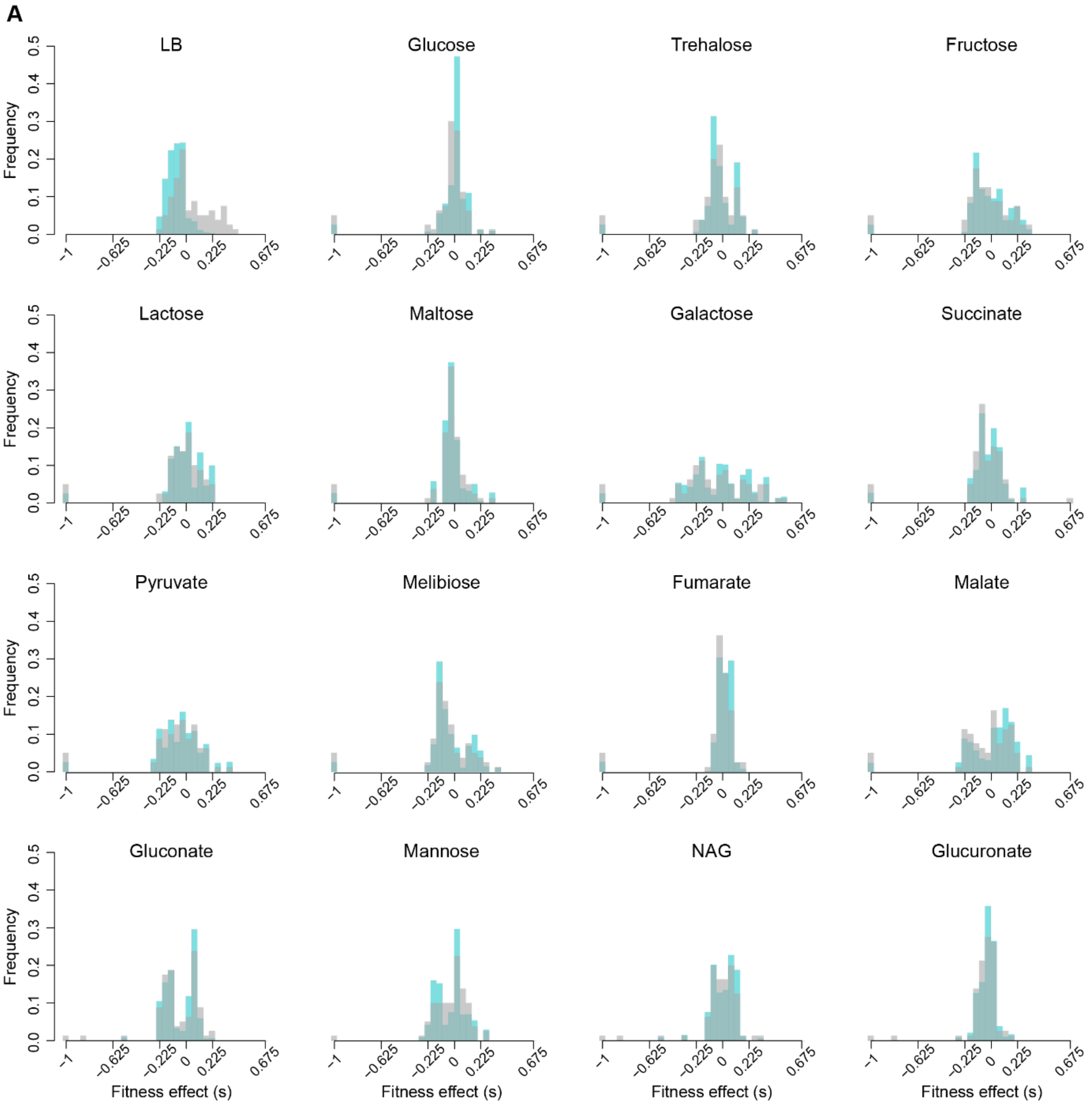

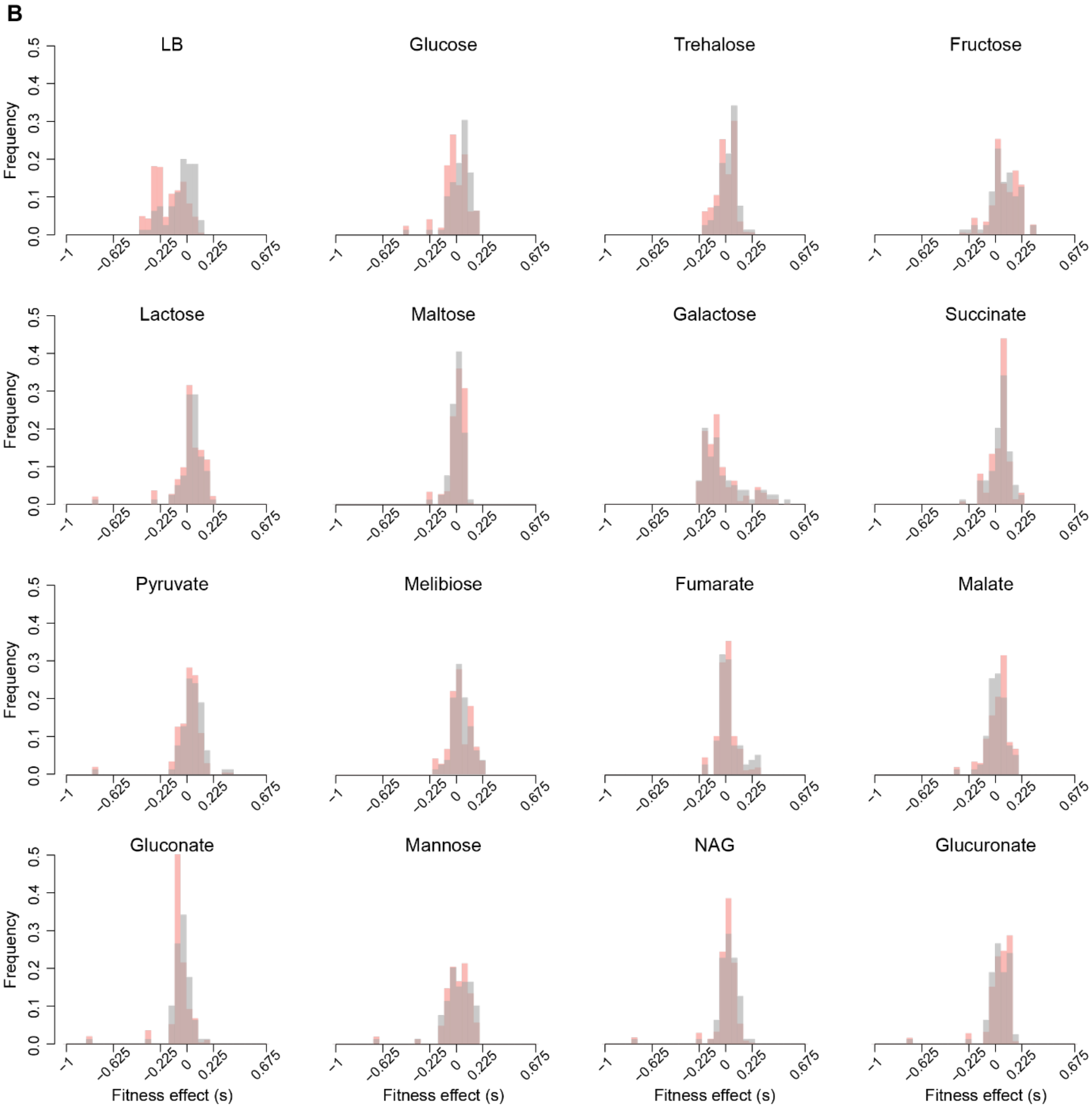
B
LB
Glucose
Trehalose
Fructose
Lactose
Maltose
Galactose
Succinate
Pyruvate
Melibiose
Fumarate
Malate
Gluconate
Mannose
NAG
Glucuronate
Frequency
0.0
0.1
0.2
0.3
0.4
0.5
−1
−0.625
−0.225
0
0.225
0.675
Fitness effect (s)

331

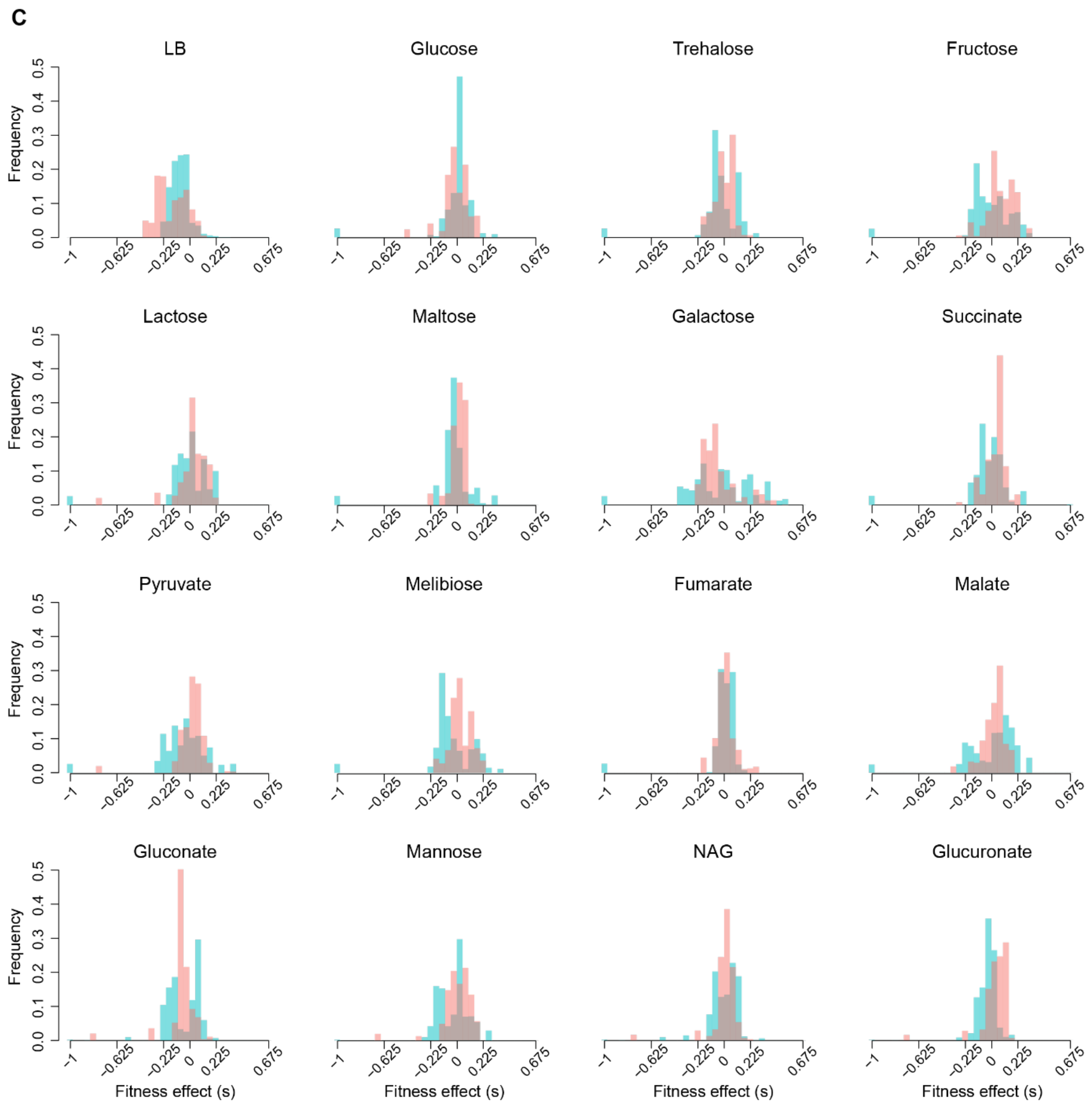
C
LB
Glucose
Trehalose
Fructose
Lactose
Maltose
Galactose
Succinate
Pyruvate
Melibiose
Fumarate
Malate
Gluconate
Mannose
NAG
Glucuronate
Frequency
0.0
0.1
0.2
0.3
0.4
0.5
−1
−0.625
−0.225
0
0.225
0.675
Fitness effect (s)

**Figure S2. Fitness consequences and incidence of pleiotropy among new mutations.** (A) Mean and (B) Median fitness effect of new mutations in WT and mutator in 16 environments. (C-F) Stacked bar plots show the proportion of mutants showing various categories of fitness effects in (C) WT, (D) Mutator, (E) WT DFE shifted towards beneficial mutations and (F) WT DFE shifted towards deleterious mutations. Each bar represents pooled data across all pairwise resource comparisons for a focal environment, except LB (panels C, E and F: *n* = 80 mutants x 14 environment pairs involving the focal environment = 1120 data points per bar; panel D: *n* = 79 mutants x 14 environment pairs involving a focal environment = 1106 data points per bar). Asterisks indicate significant differences in the proportion of fitness effects falling into each category, compared to WT (cyan asterisks) or mutator (pink asterisks) ($p < 0.05$, chi-squared tests; see Table S6).

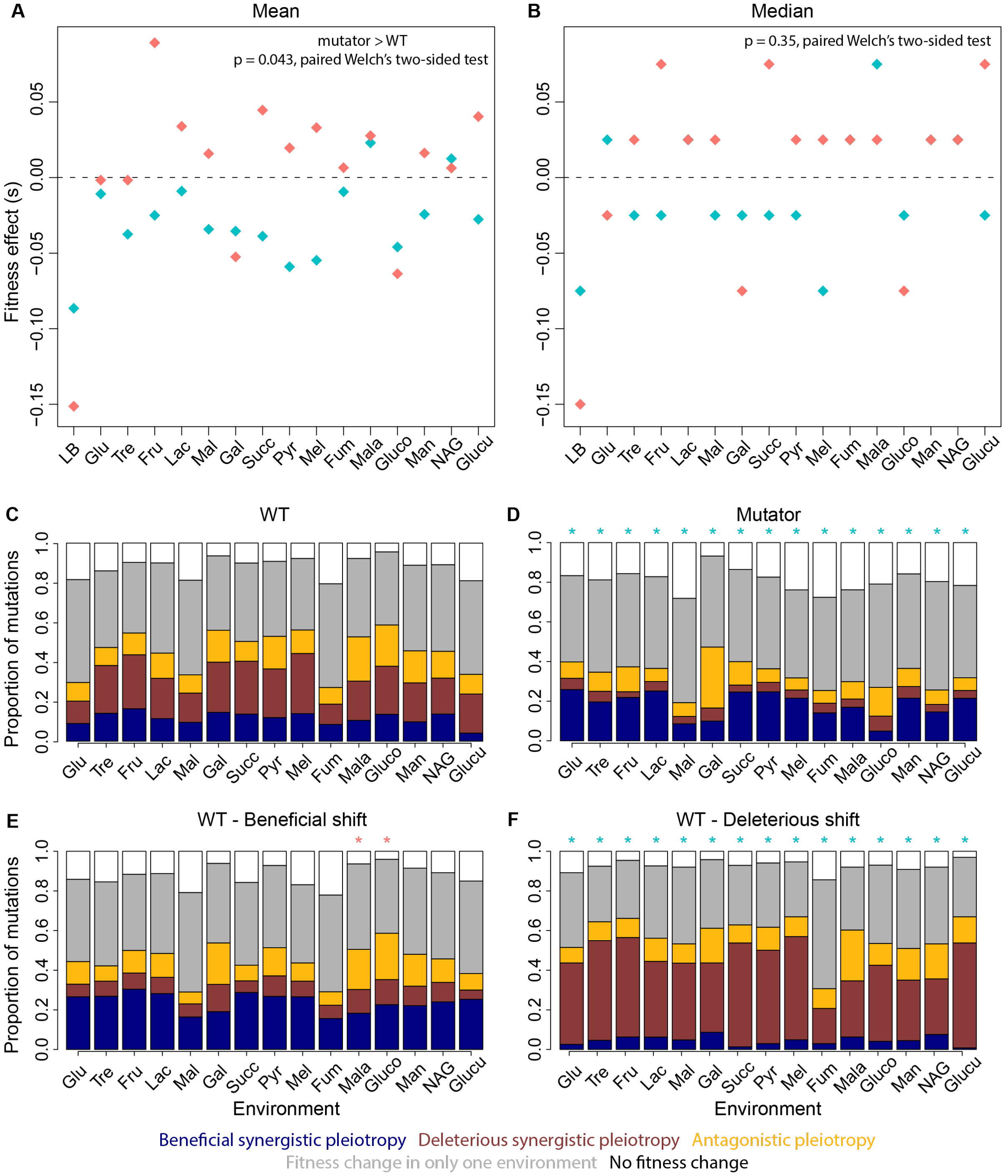

**Figure S3. Fitness effects of mutations across environments.** Heatmaps show fitness impacts (see colour key) of new mutations in (A) WT and (B) mutator for each mutation (y-axis; numbered as in Supplementary Data) across multiple environments (x-axis). Asterisks indicate mutations whose effects was tested in both WT and mutator backgrounds (Fig. 3B).

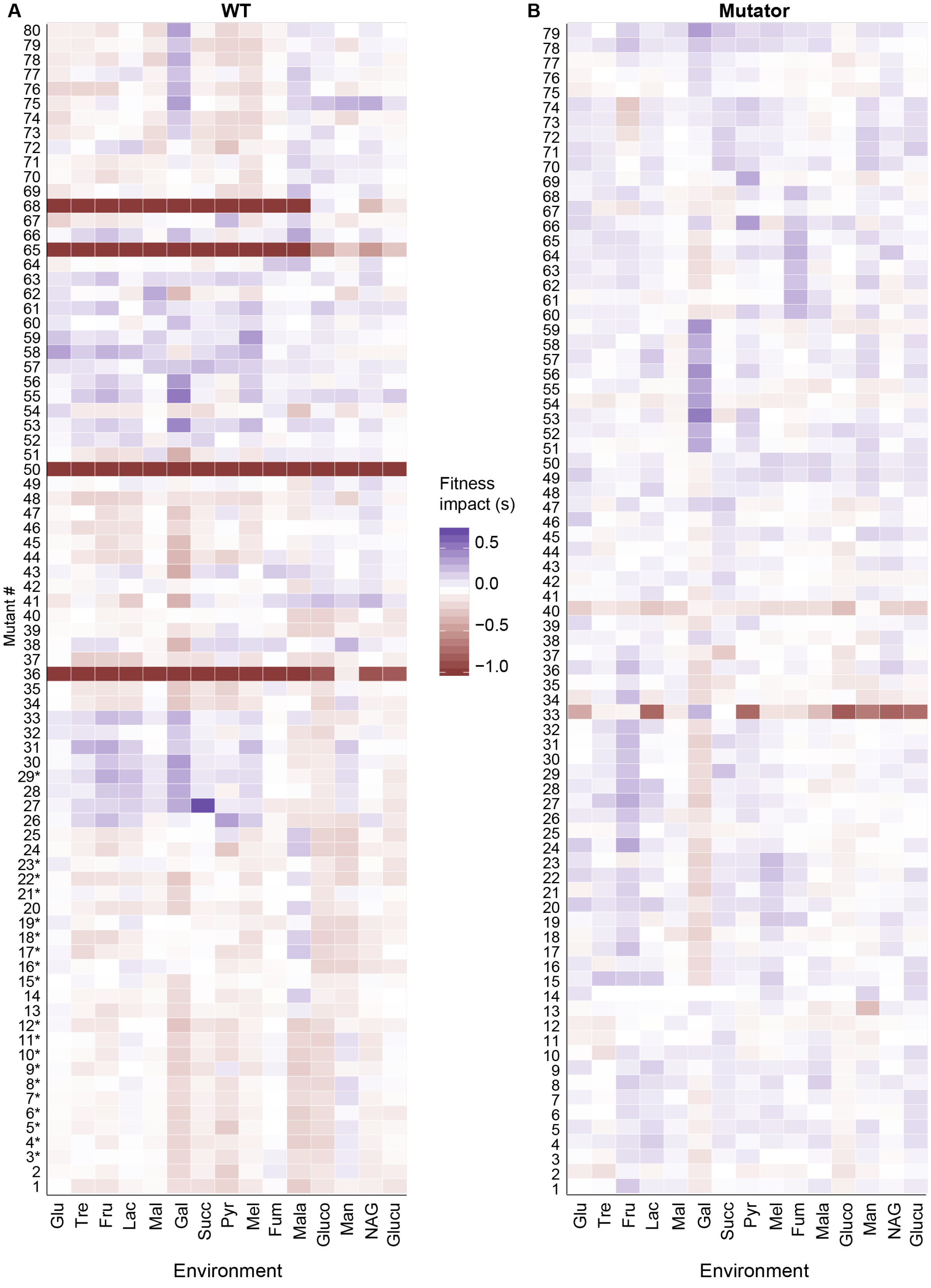

**Figure S4. BPS/Indel, coding/non-coding and synonymous/non-synonymous mutation biases are not significantly associated with fitness differences across WT and mutator.** Fitness impacts of different mutation classes, for MA-evolved isolates carrying single mutations. Boxplots (colored by strain (blue: WT, pink: mutator) and mutation class) show median, quartiles and range of fitness impacts of mutations (open diamonds: outliers). Each boxplot was constructed with n x 16 data points (n = number of mutations, shown below each plot; pooled fitness effects from 16 environments). Black horizontal lines: s = 0; black asterisks: significant differences across mutation classes within a strain (Table S9); ns: no significant difference; nd: not determined. For most comparisons, we determined significance using ANOVA (Table S9). Where sample sizes were imbalanced, we used permutation tests, subsampling 100 times from the group with the larger sample size followed by non-parametric Wilcoxon's rank sum tests. Asterisks indicate cases where >90% of permutations showed a significant difference across groups. Blue asterisks: significant differences within a mutation class across strains (Wilcoxon's rank sum test, BPS: W = 548606, p < 2.2E-16; coding mutations: W = 424210, p < 2.2E-16; non-coding mutations: W = 14150, p = 1.486E-09; non-synonymous mutations: W = 211800, p = 1.76E-09; synonymous mutations: W = 24463, p = 9.55E-07).

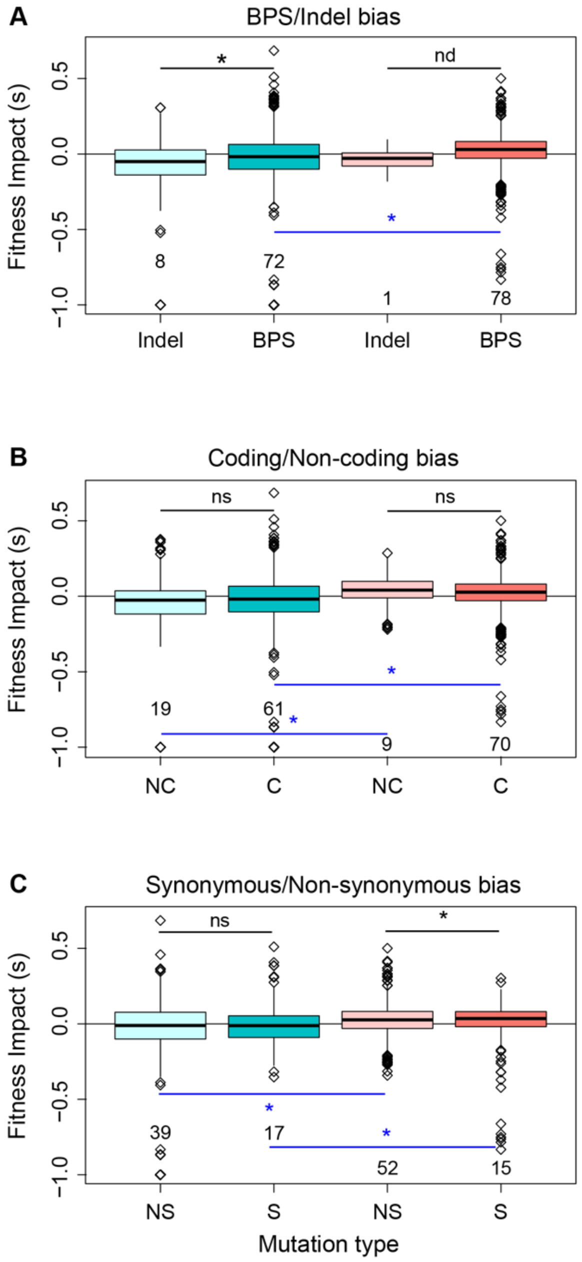

**Figure S5. Genomic aspects of mutations do not differ across strains.** Plots show the distribution of mutations in WT and mutator, with respect to various genomic factors that may influence their fitness effects. Plots are coloured by strain background (blue: WT, pink: mutator). (A) Histograms show the distribution of distance from the *E. coli* chromosomal origin of replication (*oriC*) for synonymous, nonsynonymous, and non-coding mutations. Stacked bar plots show the proportion of mutations (B) on each replichore of the genome and (C) on each DNA strand (plus vs. minus). The outcome of pairwise tests comparing the effect of strain background is noted in parentheses in panel headers (A: KS-tests, Table S10; B-C: chi-squared tests, Table S11; * = significant difference; ns = no difference).

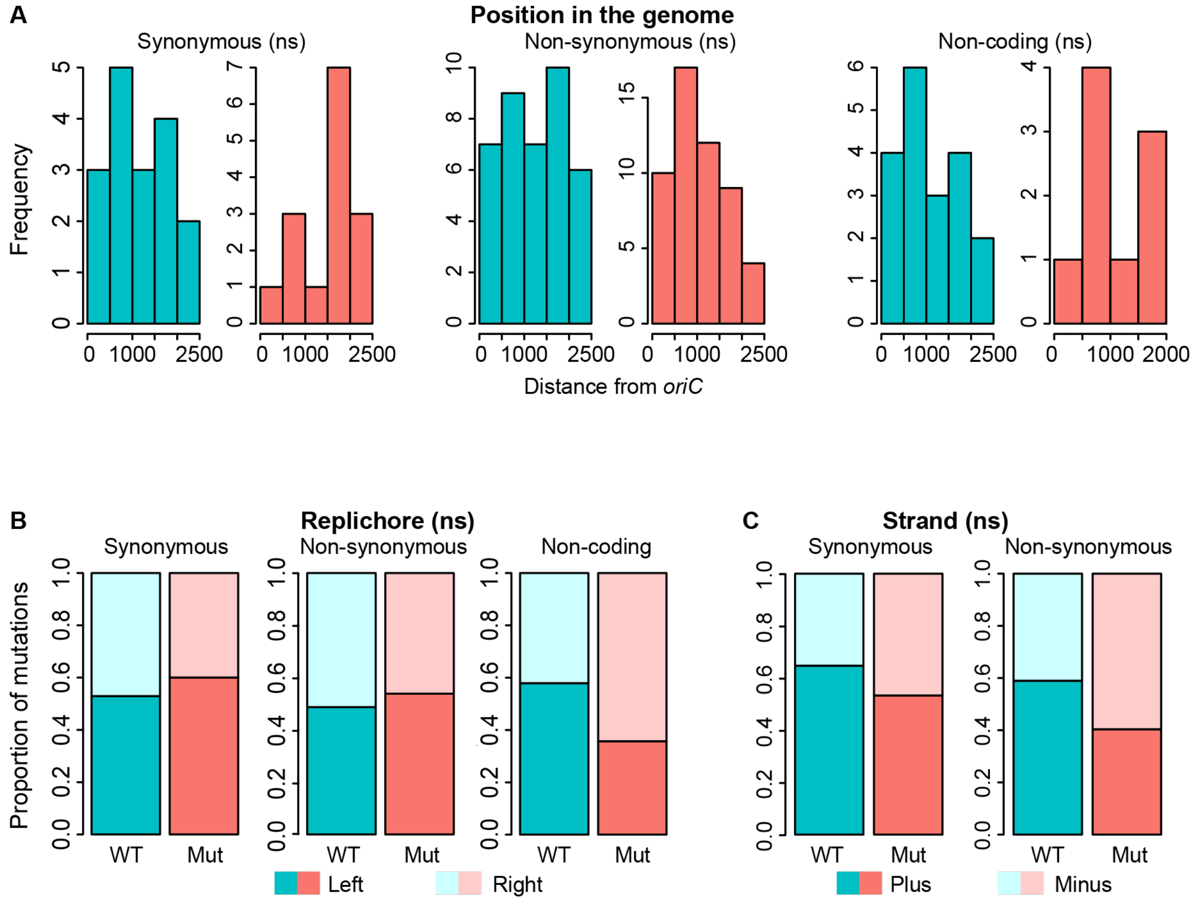

**Figure S6. Genic aspects of mutations do not differ across strains.** Plots show the distribution of mutations in WT and mutator, with respect to various genic factors that may influence their fitness effects. Plots are coloured by strain background (blue: WT, pink: mutator). (A) Histograms show the distribution of distance from the start codon for synonymous and nonsynonymous mutations. (B-F) Stacked bar plots show the proportion of mutations that are (B) of the core vs. the accessory genome of *E. coli*, (C) essential vs. non-essential for growth in LB, (D) essential vs. non-essential for growth in M9 minimal medium + glucose, (E) classified into functional GO-categories and (F) conservative vs. non-conservative amino acid changes. The outcome of pairwise tests comparing the effect of strain background is noted in parentheses in panel headers (A: KS-tests, Table S10; B-F: chi-squared tests, Table S11; * = significant difference; ns = no difference).

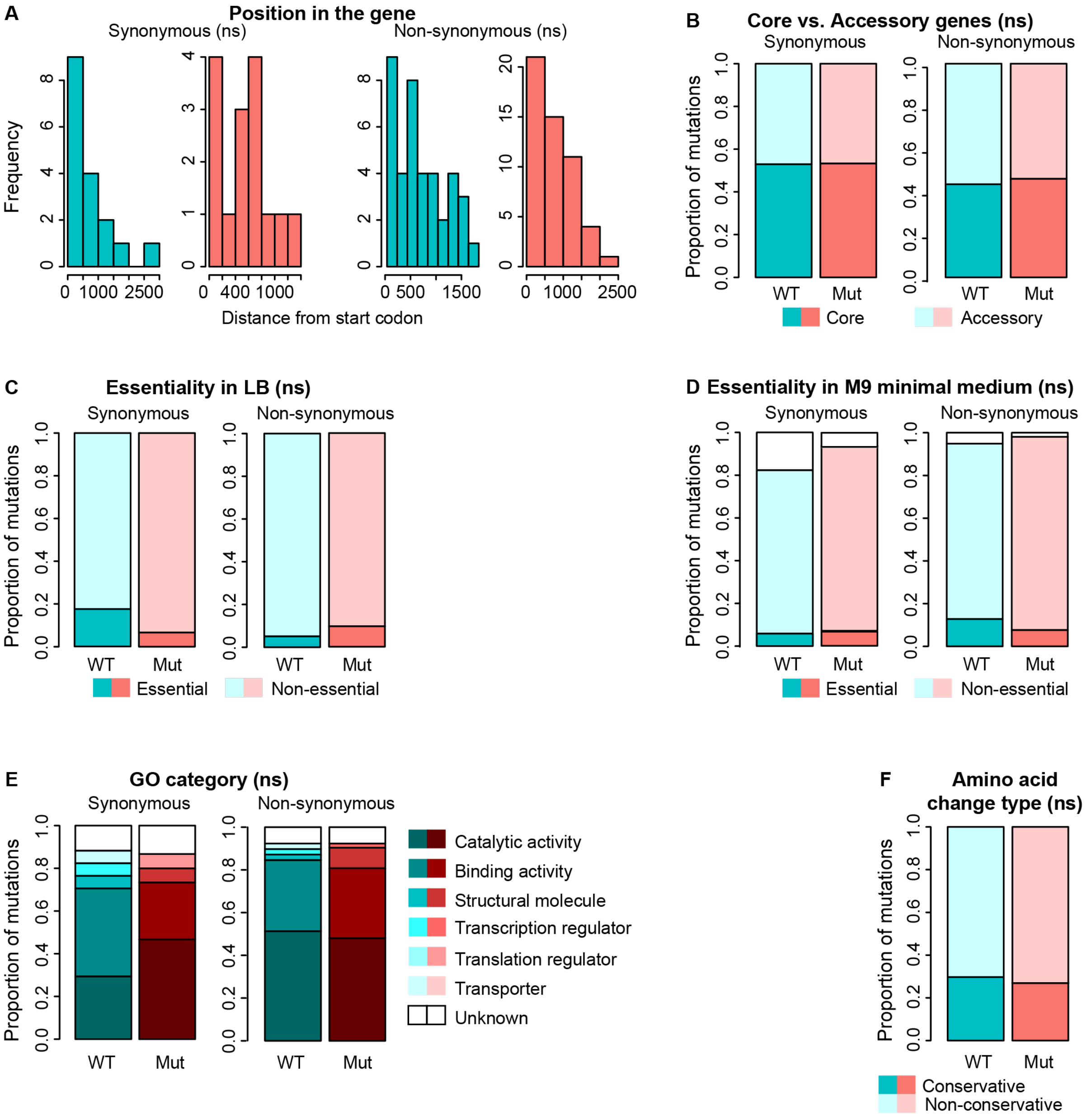

**Figure S7. Variation in simulation results.** (A) The mean and SD (coloured error bars) over 500 adaptive walks for the beneficial fraction of the DFE (open circles, offset for clarity) and mean selective effect of beneficial mutations (crosses) for mutators, versus the Tv fraction of the mutator. The wildtype has a fixed Tv fraction of 0.45 and a beneficial fraction of 0.04. While substantial walk-to-walk variability is observed, the standard error of the mean (black error bars, sometimes within symbols) indicates that uncertainty in the mean value across walks is negligible. (B) Analogous results for the deleterious fraction of the DFE (open circles) and mean deleterious effect (crosses). Black error bars indicating standard error of the mean fall well within symbol heights. (C) The difference in beneficial fractions between transitions and transversions after unbiased walks. Adaptive walks starting at a random sequence with N=200 were simulated across epistatic landscapes (K=1) with an unbiased transversion ratio of 2/3. After 50,000 steps in the simulation (mean remaining beneficial fraction 0.173), the beneficial fraction was computed for the DFE of all transitions ($f_{b\ Ts}$), and for the DFE of all transversions ($f_{b\ Tv}$). The histogram plots the percent difference in these fractions, $100(f_{b\ Tv} - f_{b\ Ts})/f_{b\ Ts}$, for 500 individual walks (each across a new randomly-generated landscape). Although on average the beneficial fraction of transitions and transversions, after these unbiased walks, is not significantly different (paired t-test, p=0.72), these results indicate that after a particular walk, either transitions or transversions may be substantially favoured.

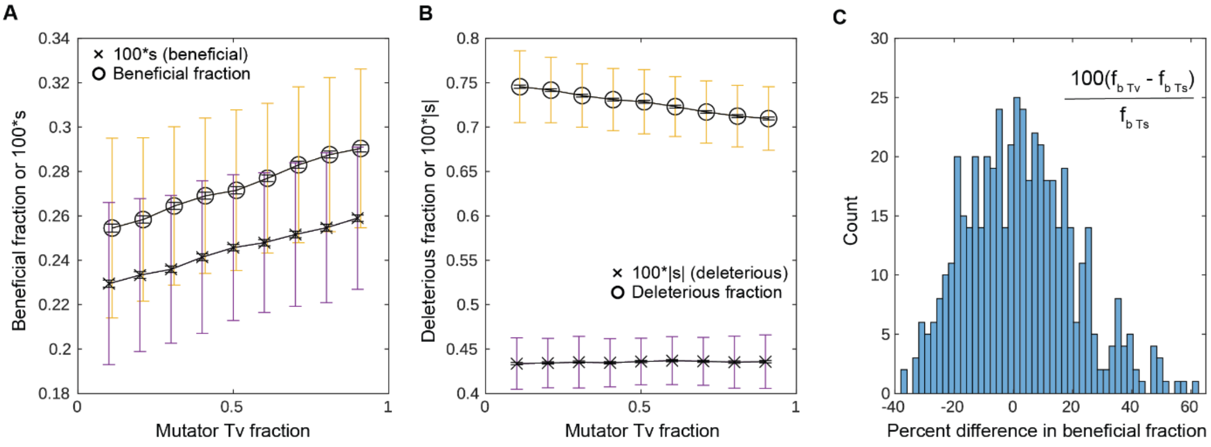

**Figure S8. Waterfall plots showing the impact of altering Tv bias at the end of an adaptive walk.** Adaptive walks as described for Fig. 4 were simulated with a WT (ancestor) Tv bias fixed to a single value between 0.1 and 0.9; walks at this fixed bias were simulated until the WT was well-adapted ($f_b$ = 0.04). At the end of these walks, the bias was shifted successively from 0.1 to 0.9 by increments of 0.1. At each shifted bias value, a new DFE was generated. (A) $f_b$ increases when the shift in bias reverses the WT bias, and decreases when the shift reinforces the WT bias. (B) The same trend is observed for $s_b$. (C) $f_d$ decreases when the bias is reversed and increases when the bias is reinforced. (D) Changes is $s_d$ are small relative to changes in $s_b$, but $s_d$ increases slightly when the bias is reversed.

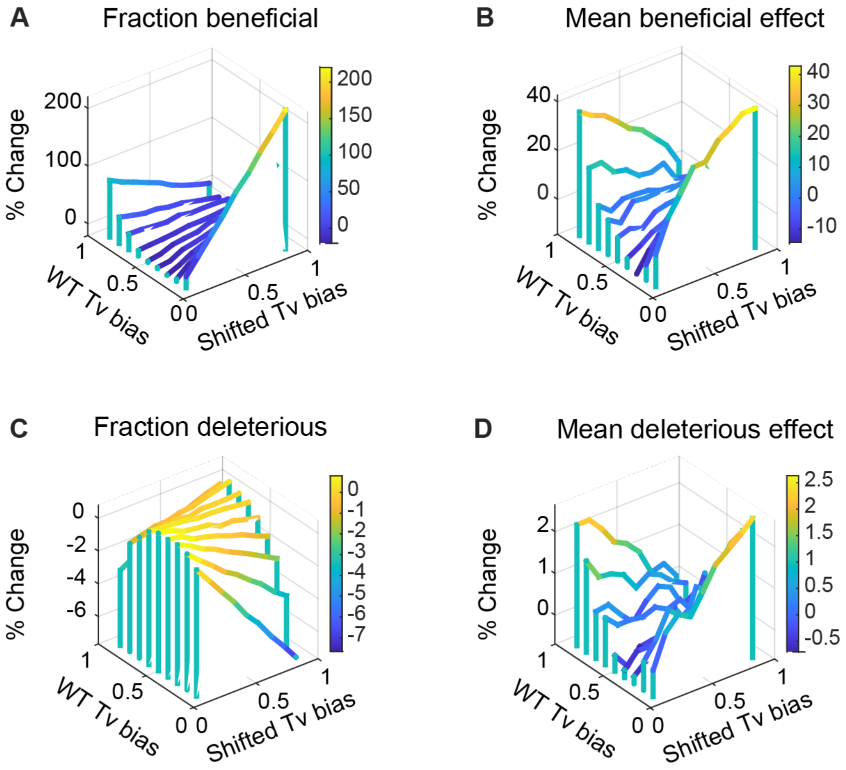

**Figure S9. Adaptive walk simulations when altering GC→AT bias.** Adaptive walks were simulated starting with a random ancestor sequence (50% GC content). The proportion of GC→CG and AT→TA mutations was fixed at 10%, mimicking our WT. (A) Change in mean population fitness, fraction of beneficial and deleterious mutations ($f_b$ and $f_d$), and magnitude of beneficial and deleterious effects ($s_b$ and $s_d$) over the course of adaptive walks. The GC→AT bias = GC→AT/(GC→AT+AT→GC) was fixed at 62%. Average GC content at the end of the walk was 35.2%. (B) Impact of altering GC→AT bias on $f_b$, at different points along the adaptive walk. Here, the walk was paused, the sequence was fixed (and thus the GC content was fixed), but new DFEs were computed using different values of the GC→AT bias. (C–F) Waterfall plots showing the impact of altering GC→AT mutation bias (% change, Mutator – WT), as a function of WT (ancestor) and mutator (new) bias, for a well-adapted WT population ($f_b$ = 0.04).

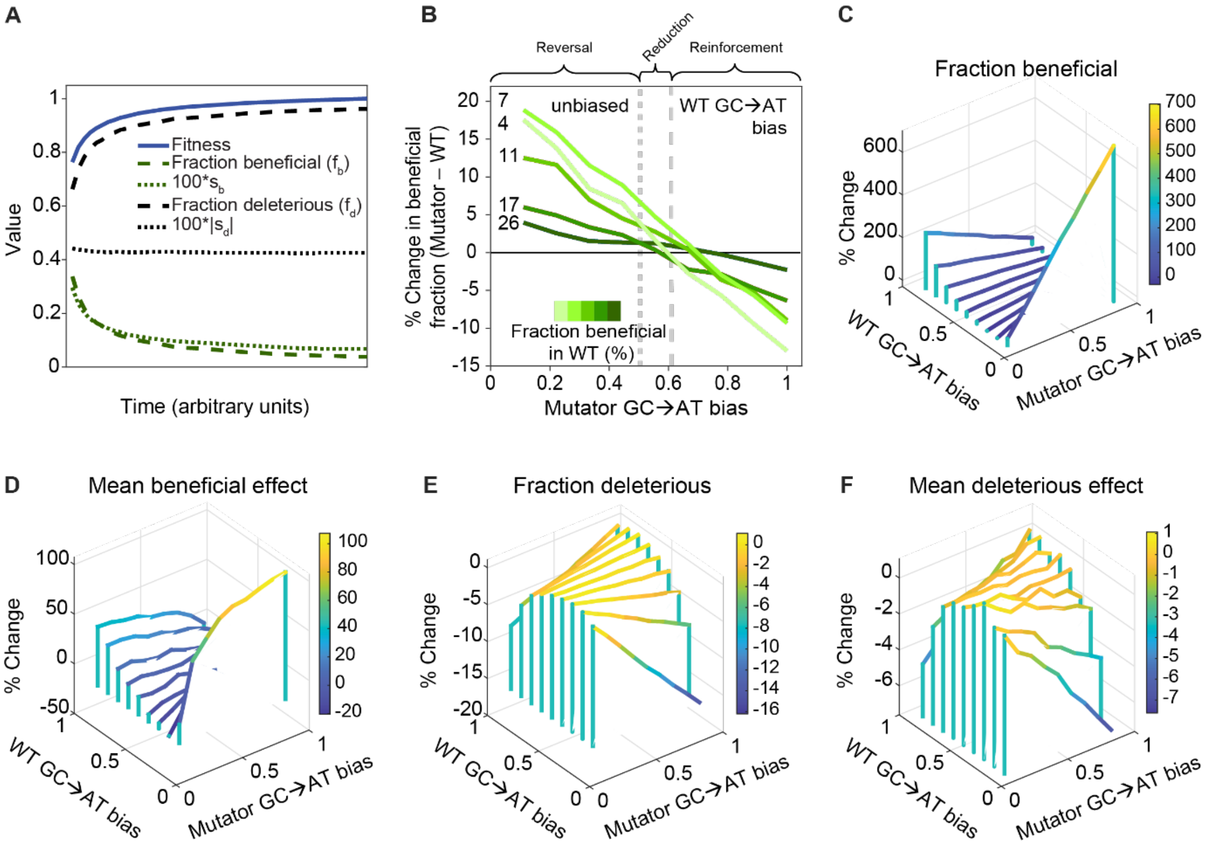

**Figure S10. Epistasis and degeneracy in adaptive walk simulations.** Adaptive walks were simulated for 10000, 100 or 15 timesteps (squares) and for 35000, 500 or 70 timesteps (circles) for N = 200, 20 or 8 respectively, for a WT Tv bias of 0.45. The degree of epistasis and degeneracy in the fitness landscape increases with increasing values of the fraction K/N, and N=8, K=8 implies a completely uncorrelated landscape. (A) The mean over 500 adaptive walks for the beneficial fraction of the DFE ($f_b$) of the WT, as K changes from 1 to 8 for a given N. (B) The percent increase in $f_b$ at the end of each walk for a bias-shifted strain with Tv bias 0.9 (relative to WT). The benefit of the shift in mutational bias holds across a wide range of N and K values, and for wildtype beneficial fractions of 5 to 35%. (C) Distribution of beneficial mutations sampled during the adaptive walks. Adaptive walks with N=200 and K=1 were simulated from a random sequence with Tv bias = 0.45 until $f_b$ reached 35%. The sequence of this "ancestor" was saved, and the walk was continued until $f_b$ reached 25%. This second adaptive walk was repeated 500 times, starting with the same ancestor for the same number of timesteps. Comparing the 500 evolved sequences with the ancestor, we found 343 distinct substitutions. The plot shows a histogram of the fraction of genomes sharing each of these unique substitutions; 25.9% are shared by less than 1% of the evolved lines, and 69.4% are shared by less than 10% of the evolved lines. The second set of 500 walks was then repeated, starting with the same ancestor, but with Tv bias = 0.9 (bias-shifted strain). In this case 345 distinct substitutions were observed; 32.4% are shared by less than 1% of the evolved lines, and 73.0% are shared by less than 10% of the evolved lines. Thus, adaptive trajectories in our simulation studies do not exhaust available beneficial mutations, but make a wide diversity of substitutions with little parallel evolution.

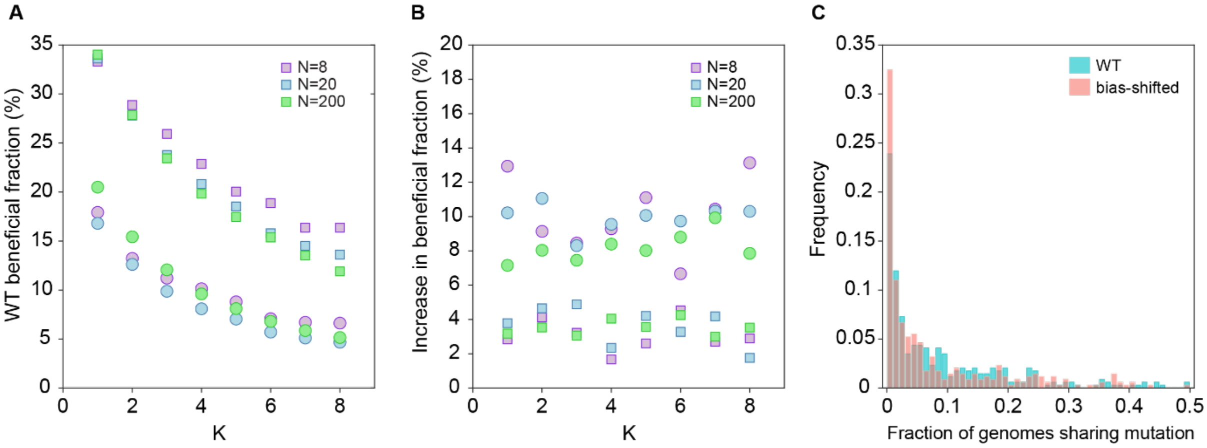

**Figure S11. Results of full population simulations with multiple segregating mutations.** Populations were evolved at mutation rates of $10^{-6}$, $10^{-5}$ and $10^{-4}$ mutations per genome per generation, corresponding to averages of 1.9, 9.7 and 77.8 segregating genotypes in the population at any time (results for $10^{-4}$ are shown in the main text). (A and C) Population mean fitness increases over time (means of 1000 independent replicate populations are illustrated). The beneficial fraction and mean beneficial effect size decrease as evolution progresses, while the deleterious fraction increases. (B and D) As described for the adaptive walk simulations, we pause the population at various time points and find the most common genotype in the population. We then temporarily shift the Tv bias to a new value, and create 500 random single-step mutants from the most common genotype using this shifted probability of Tv to generate the mutations. Lines are labelled with the beneficial fraction of the DFE in the ancestor at the time evolution was paused. We then plot the percent change in this value for shifted Tv biases between 0.1 and 0.9. Because fb decreases over time as the ancestor evolves, lighter lines indicate later time points in adaptation. The main effect we observe here is that as the mutation rate increases, the population adapts more rapidly and thus these trends emerge more quickly (compare x-axis ranges across panels A and C). In other words, increasing the overall mutational supply does not diminish the effects of bias shifts, but accelerates their emergence as the population rapidly evolves.

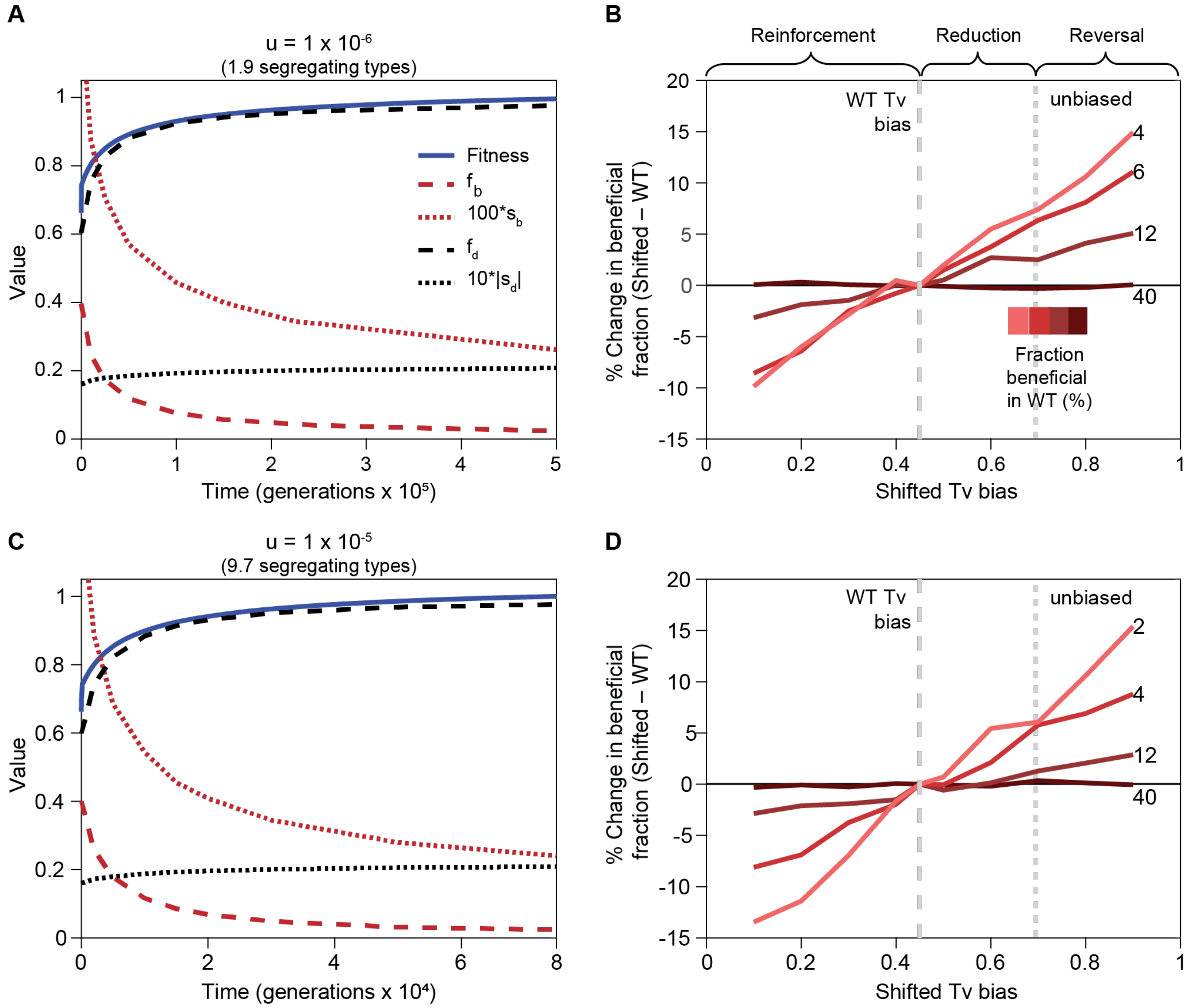

**Figure S12. Results of full population simulations using a codon-based fitness model.** Genotypes are of n nucleotides and mutations occur at the nucleotide level. In this model, nucleotides were translated to amino acids before computing fitness. Fitness was computed using an NK-landscape that depended only on amino acid identity, with N = n/3. Thus, synonymous mutations are neutral in this fitness landscape. (A) Change in mean population fitness, fraction of beneficial and deleterious mutations ($f_b$ and $f_d$), and magnitude of beneficial and deleterious effects ($s_b$ and $s_d$) over the course of adaptive walks of a WT ancestor. The fraction of neutral mutations is also shown. (B) Impact of altering Tv bias on $f_b$, at different points along the adaptive walk. We note that increasing the Tv bias reduces the neutral fraction due to the redundancy of the genetic code, and thus in the simulations both $f_b$ and $f_d$ increase as Tv bias increases. To correct for this effect, we plot the change in the beneficial fraction among non-neutral mutations, that is, the percent change in $f_b/(f_b + f_d)$. We also note that in the experimental data, the fraction $f_b/(f_b + f_d)$ is higher in the mutator than in the WT in 11 of 16 environments. Parameter values: n=99, K=3, $10^{-4}$ mutations per genome per generation, means of 1000 independent replicate populations are shown.

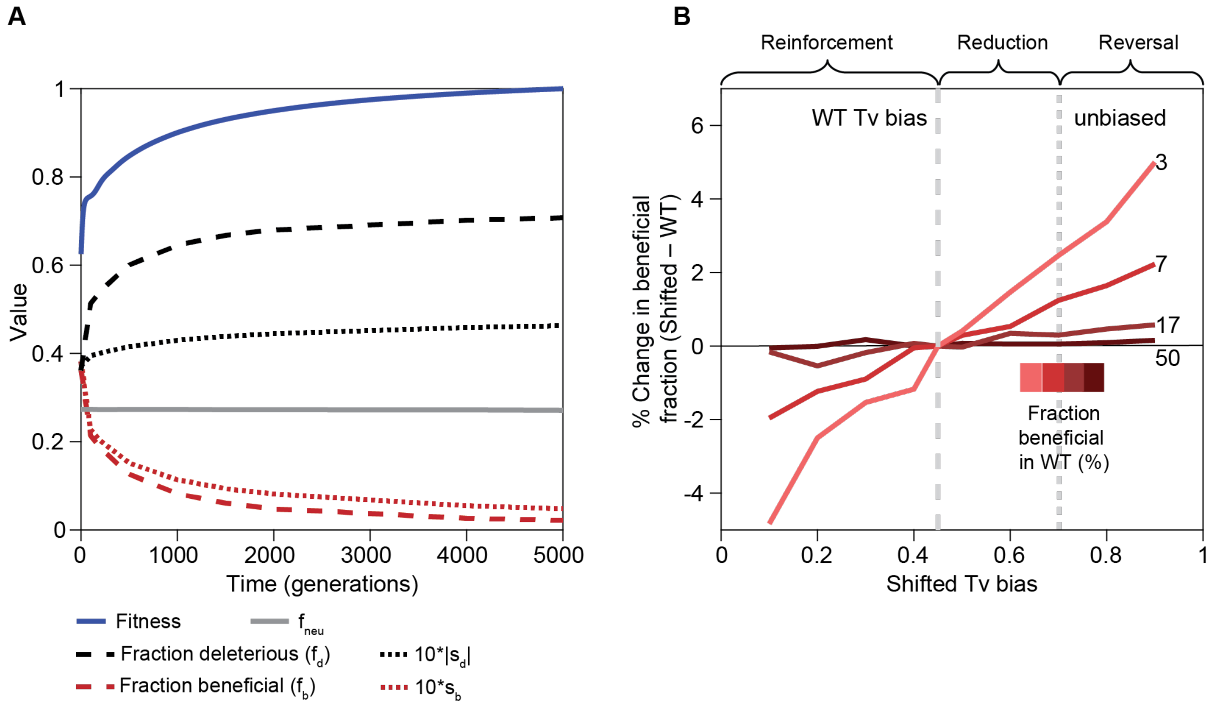

**Figure S13. Phylogenomic analysis of additional DNA repair genes whose effect on the mutation spectrum is not known.**

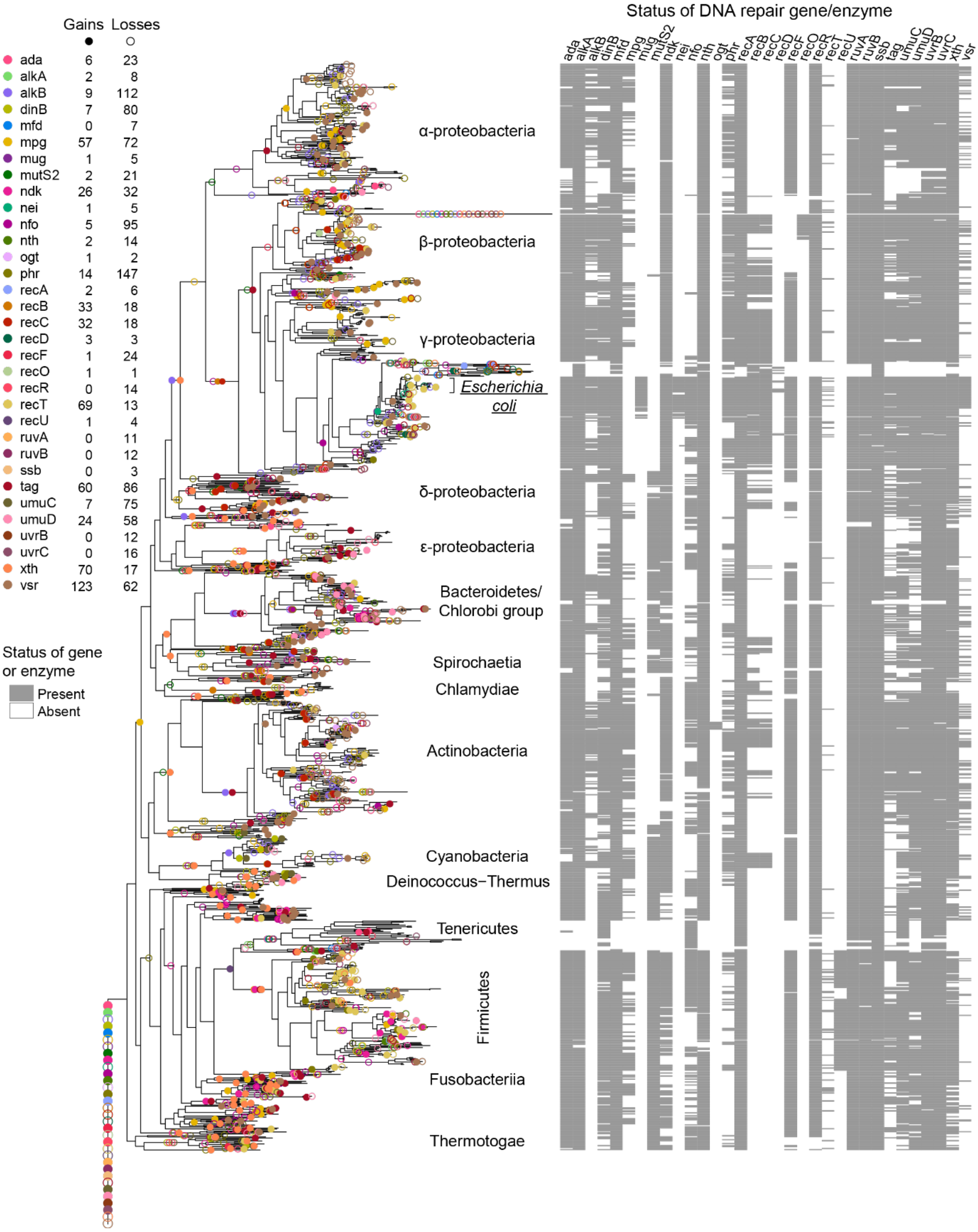

**Figure S14. Inference of changes in the direction of mutation bias in the bacterial phylogeny.** (A) Number of lineages showing different types of bias shifts expected by chance alone. Error bars represent variation across 10,000 simulations. (B–C) Frequency distributions of occurrence of successive bias shifts in the same or opposite direction per lineage, across 10,000 simulations for (B) Tv bias and (C) GC→AT bias. Red lines indicate the observed number of successive bias shifts in the same or opposite direction per lineage (shown in Fig. 5C); black lines indicate the median of the distribution of simulation results (“expected”).

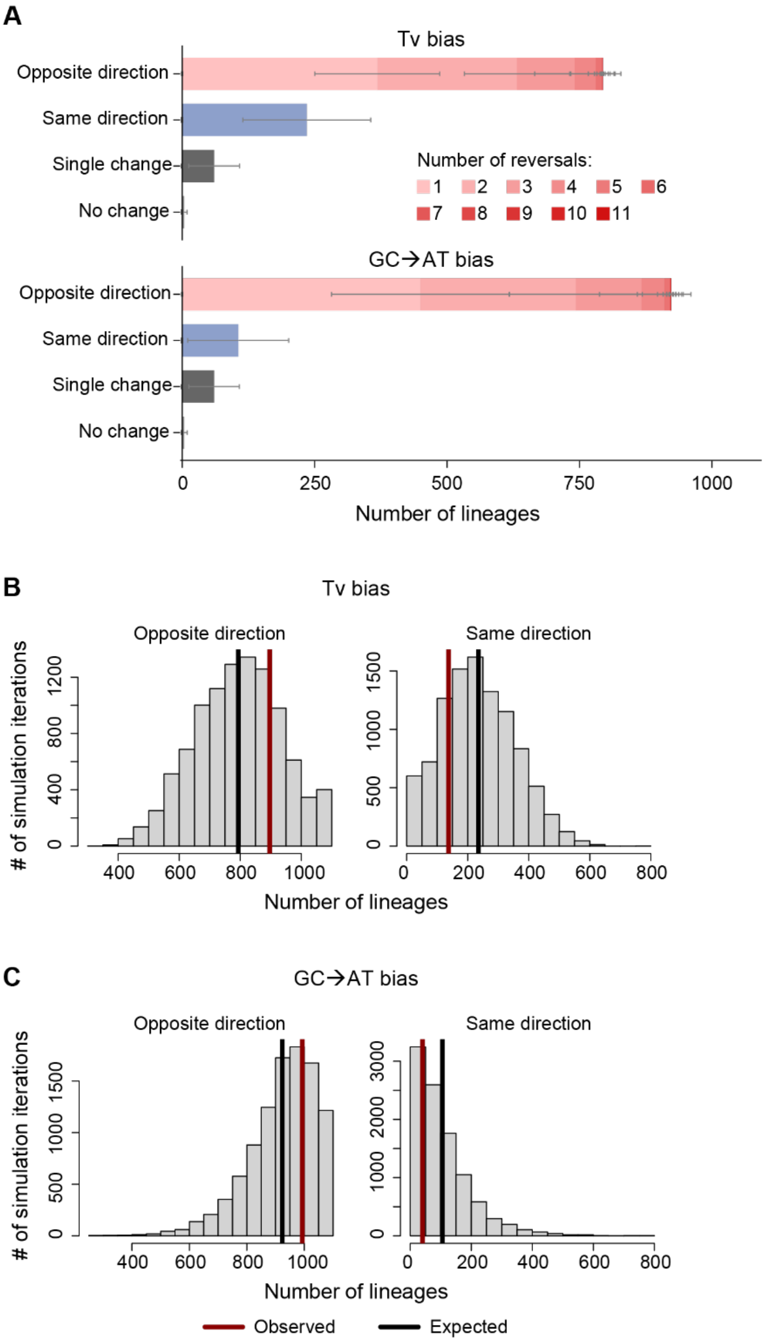

### SUPPLEMENTARY TABLES

**Table S1. Transversion (Tv) bias across different "wild type" microbial species, from mutation accumulation studies.** Tv bias = Number of Tv mutations / (Number of Tv + Number of Ts mutations). GC→AT bias = Number of GC→AT mutations/(Number of GC→AT mutations + Number of AT→GC mutations). Larger values indicate more transversions or GC→AT mutations.

| Species | Tv bias | GC→AT bias | Reference |
|---|---|---|---|
| *Bacillus subtilis* | 0.249 | 0.494 | (19) |
| *Vibrio fischeri* | 0.539 | 0.737 | (20) |
| *Vibrio cholerae* | 0.391 | 0.625 | (20) |
| *Vibrio cholerae* | 0.367 | 0.607 | (21) |
| *Pseudomonas fluorescens* | 0.368 | 0.643 | (22) |
| *Pseudomonas aeruginosa* | 0.34 | 0.750 | (23) |
| *Deinococcus radiodurans* | 0.37 | 0.528 | (24) |
| *Mesoplasma florum* | 0.452 | 0.855 | (25) |
| *Mycobacterium smegmatis* | 0.403 | 0.597 | (26) |
| *Mycobacterium smegmatis* | 0.377 | 0.714 | (27) |
| *Corynebacterium glutamicum* | 0.3 | 0.604 | (28) |
| *Saccharomyces cerevisiae* | 0.62 | 0.500 | (29) |
| *Schizoaccharomyces pombe* | 0.563 | 0.610 | (30) |

**Table S2: Output of chi-square tests comparing the proportion of beneficial, neutral, and deleterious mutations, or beneficial vs. all other mutations across WT and mutator in each environment.**

| Environment | Beneficial, neutral, deleterious | | Beneficial vs. others | |
|---|---|---|---|---|
| | **Statistic** | **p** | **Statistic** | **p** |
| LB | 0.90 | 0.639 | 1.65E-29 | 1 |
| Glucose | 6.75 | **0.039** | 4.71E-31 | 1 |
| Trehalose | 8.87 | **0.019** | 4.20 | **0.040** |
| Fructose | 25.85 | **1.30E-05** | 20.60 | **5.67E-06** |
| Lactose | 6.74 | **0.039** | 5.15 | **0.023** |
| Maltose | 13.05 | **0.003** | 11.02 | **0.001** |
| Galactose | 8.05 | **0.024** | 6.04 | **0.014** |
| Succinate | 24.66 | **1.77E-05** | 22.40 | **2.21E-06** |
| Pyruvate | 14.18 | **0.002** | 8.94 | **0.003** |
| Melibiose | 36.49 | **9.54E-08** | 15.98 | **6.41E-05** |
| Fumarate | 3.04 | 0.233 | 1.26 | 0.261 |
| Malate | 10.53 | **0.009** | 0.70 | 0.403 |
| Gluconate | 20.08 | **1.16E-04** | 12.64 | **3.78E-04** |
| Mannose | 8.21 | **0.024** | 7.10 | **0.008** |
| NAG | 23.81 | **2.16E-05** | 1.70 | 0.192 |
| Glucuronate | 45.94 | **1.70E-09** | 36.08 | **1.89E-09** |

**Table S3. Numbers of mutations of different types observed in this study and a previous study** (4), **which measured mutation rates and spectra in MA lines of a Δ*mutY* genotype of *E. coli*.** Higher bias values indicate more Tv or GC→AT mutations. Chi-square tests compare mutation spectra across strains, and between this study and the previous study.

| | **This study** | | **Comparison** | | | | | |
|---|---|---|---|---|---|---|---|---|
| | **WT** | **Δ*mutY*** | **WT (this study vs. previous study)** | | **Δ*mutY* (this study vs. previous study)** | | **WT vs. Δ*mutY* (this study)** | |
| **Mutation rate x $10^{-11}$ ($bp^{-1}$ $gen^{-1}$)** | 9.3 | 89 | | | | | | |
| **Mutation type** | | | **$\chi^2$** | **p** | **$\chi^2$** | **p** | **$\chi^2$** | **p** |
| BPS | 237 | 669 | | | | | 34.2 | **4.8E-09** |
| Indel | 25 | 9 | | | | | | |
| Indel bias | 0.095 | 0.013 | | | | | | |
| Coding | 167 | 577 | 2.13 | 0.14 | 1.46 | 0.23 | 28.6 | **8.8E-08** |
| Noncoding | 70 | 92 | | | | | | |
| Noncoding bias | 0.295 | 0.137 | | | | | | |
| Synonymous | 60 | 162 | 0.83 | 0.36 | 0.25 | 0.62 | 3.45 | 0.063 |
| Non-synonymous | 107 | 415 | | | | | | |
| Non-synonymous bias | 0.64 | 0.719 | | | | | | |
| Transition | 131 | 72 | 0.01 | 0.91 | 10.7 | **0.001** | 196.8 | **1.0E-44** |
| Transversion | 106 | 597 | | | | | | |
| Tv bias | 0.447 | 0.892 | | | | | | |
| AT→GC | 86 | 22 | 0.26 | 0.61 | 3.44 | 0.064 | 200.7 | **1.4E-45** |
| GC→AT | 125 | 632 | | | | | | |
| No change | 26 | 15 | | | | | | |
| GC→AT bias | 0.592 | 0.966 | | | | | | |

**Table S4. Estimated supply of beneficial mutations in WT and mutator.** The genome size of *E. coli* K-12 MG1655 is 4641650 bp. μ = mutation rate $bp^{-1}$ $generation^{-1}$; $f_b$ = fraction beneficial mutations (Fig. 1); $S_b$ = supply of beneficial mutations $genome^{-1}$ $generation^{-1}$ (μ x $f_b$).

| **Environment** | **WT (μ = 9.3 x $10^{-11}$)** | | **Δ*mutY* (μ = 8.9 x $10^{-10}$)** | | | **Fold increase in Δ*mutY*** | |
|---|---|---|---|---|---|---|---|
| | **$f_b$** | **$S_b$** | **$f_b$** | **$S_b$** | **$S_{b(WT\ DFE)}$** | **$S_b$ (Δ*mutY*)/ $S_b$ (WT)** | **$S_{b(WT\ DFE)}$ (Δ*mutY*)/ $S_b$ (WT)** |
| LB | 0.06 | 0.00002 | 0.05 | 0.0002 | 0.0002 | 9.02 | 9.57 |
| Glucose | 0.22 | 0.00010 | 0.34 | 0.0014 | 0.0009 | 14.37 | 9.57 |
| Trehalose | 0.28 | 0.00012 | 0.35 | 0.0014 | 0.0011 | 12.14 | 9.57 |
| Fructose | 0.35 | 0.00015 | 0.58 | 0.0024 | 0.0014 | 15.69 | 9.57 |
| Lactose | 0.32 | 0.00014 | 0.43 | 0.0018 | 0.0013 | 12.92 | 9.57 |
| Maltose | 0.15 | 0.00007 | 0.31 | 0.0013 | 0.0006 | 19.32 | 9.57 |
| Galactose | 0.36 | 0.00015 | 0.20 | 0.0008 | 0.0015 | 5.36 | 9.57 |
| Succinate | 0.25 | 0.00011 | 0.60 | 0.0024 | 0.0010 | 22.66 | 9.57 |
| Pyruvate | 0.28 | 0.00012 | 0.41 | 0.0017 | 0.0012 | 13.78 | 9.57 |
| Melibiose | 0.26 | 0.00011 | 0.37 | 0.0015 | 0.0011 | 13.49 | 9.57 |
| Fumarate | 0.33 | 0.00014 | 0.21 | 0.0008 | 0.0013 | 6.06 | 9.57 |
| Malate | 0.54 | 0.00023 | 0.47 | 0.0019 | 0.0022 | 8.20 | 9.57 |
| Gluconate | 0.37 | 0.00016 | 0.08 | 0.0003 | 0.0015 | 2.15 | 9.57 |
| Mannose | 0.22 | 0.00010 | 0.40 | 0.0017 | 0.0009 | 17.38 | 9.57 |
| NAG | 0.43 | 0.00019 | 0.28 | 0.0011 | 0.0018 | 6.17 | 9.57 |
| Glucuronate | 0.06 | 0.00003 | 0.54 | 0.0022 | 0.0003 | 80.35 | 9.57 |
| Median | 0.30 | 0.00012 | 0.36 | 0.0015 | 0.0011 | 13.21 | 9.57 |
| Mean | 0.31 | 0.00012 | 0.36 | 0.0014 | 0.0012 | 16.19 | 9.57 |
| Paired t-test comparing fold increase in Δ*mutY* $f_b$ calculated using its own vs. WT DFE | | | | | | | t = 1.47<br>p = 0.16 |

**Table S5. Estimated genetic load due to deleterious mutations in WT and mutator.** The genome size of *E. coli* K-12 MG1655 is 4641650 bp. μ = mutation rate $bp^{-1}$ $generation^{-1}$; $f_d$ = fraction deleterious mutations (Fig. 1); $L_d$ = Load from deleterious mutations $genome^{-1}$ $generation^{-1}$.

| **Environment** | **WT (μ = 9.3 x $10^{-11}$)** | | **Δ*mutY* (μ = 8.9 x $10^{-10}$)** | | | **Fold increase in Δ*mutY*** | |
|---|---|---|---|---|---|---|---|
| | **$f_d$** | **$L_d$** | **$f_d$** | **$L_d$** | **$L_{d(WT\ DFE)}$** | **$L_d$ (Δ*mutY*)/$L_d$ (WT)** | **$L_{d(WT\ DFE)}$ (Δ*mutY*)/ $L_d$ (WT)** |
| LB | 0.66 | 0.00028 | 0.72 | 0.00297 | 0.0027 | 10.54 | 9.57 |
| Glucose | 0.17 | 0.00007 | 0.27 | 0.00109 | 0.0007 | 14.73 | 9.57 |
| Trehalose | 0.46 | 0.00020 | 0.24 | 0.00098 | 0.0019 | 4.95 | 9.57 |
| Fructose | 0.45 | 0.00019 | 0.09 | 0.00038 | 0.0018 | 1.95 | 9.57 |
| Lactose | 0.33 | 0.00014 | 0.15 | 0.00062 | 0.0013 | 4.46 | 9.57 |
| Maltose | 0.30 | 0.00013 | 0.10 | 0.00039 | 0.0012 | 3.00 | 9.57 |
| Galactose | 0.43 | 0.00018 | 0.65 | 0.00267 | 0.0018 | 14.49 | 9.57 |
| Succinate | 0.42 | 0.00018 | 0.12 | 0.00049 | 0.0017 | 2.73 | 9.57 |
| Pyruvate | 0.46 | 0.00019 | 0.18 | 0.00073 | 0.0019 | 3.74 | 9.57 |
| Melibiose | 0.58 | 0.00025 | 0.13 | 0.00055 | 0.0024 | 2.24 | 9.57 |
| Fumarate | 0.11 | 0.00005 | 0.14 | 0.00059 | 0.0004 | 13.01 | 9.57 |
| Malate | 0.31 | 0.00013 | 0.17 | 0.00071 | 0.0013 | 5.40 | 9.57 |
| Gluconate | 0.49 | 0.00021 | 0.61 | 0.00250 | 0.0020 | 11.91 | 9.57 |
| Mannose | 0.41 | 0.00017 | 0.23 | 0.00093 | 0.0017 | 5.35 | 9.57 |
| NAG | 0.30 | 0.00013 | 0.09 | 0.00037 | 0.0012 | 2.86 | 9.57 |
| Glucuronate | 0.31 | 0.00013 | 0.08 | 0.00031 | 0.0013 | 2.32 | 9.57 |
| Median | 0.4129 | 0.0002 | 0.1631 | 0.0007 | 0.0017 | 4.71 | 9.57 |
| Mean | 0.3864 | 0.0002 | 0.2485 | 0.0010 | 0.0016 | 6.48 | 9.57 |
| Paired t-test comparing fold increase in Δ*mutY* $L_d$ calculated using its own vs. WT DFE | | | | | | | $t = -2.62$, $p = 0.019$ |

**Table S6. Chi-square tests comparing the median proportions of pleiotropic fitness impacts of mutations in WT vs. mutator; WT DFE shifted towards beneficial mutations and WT DFE shifted towards deleterious mutations; in each of 15 focal minimal medium environments.** p values are corrected for multiple comparisons using the Benjamini-Hochberg correction for false discovery rate.

| **Environments** | **WT vs. Mutator** | | **Mutator vs. WT beneficial shift** | | **WT vs. WT deleterious shift** | |
|---|---|---|---|---|---|---|
| | **Statistic** | **p** | **Statistic** | **p** | **Statistic** | **p** |
| Glucose | 31.04 | **3.75E-06** | 0.89 | 0.95 | 43.13 | **4.87E-08** |
| Trehalose | 30.42 | **4.64E-06** | 2.09 | 0.94 | 31.36 | **2.58E-06** |
| Fructose | 42.95 | **3.18E-08** | 4.96 | 0.73 | 35.03 | **6.24E-07** |
| Lactose | 41.40 | **4.75E-08** | 4.14 | 0.78 | 39.33 | **1.28E-07** |
| Maltose | 22.85 | **1.45E-04** | 2.98 | 0.93 | 38.63 | **1.56E-07** |
| Galactose | 54.35 | **6.67E-10** | 5.79 | 0.65 | 40.81 | **8.76E-08** |
| Succinate | 44.63 | **2.37E-08** | 1.87 | 0.94 | 41.19 | **8.76E-08** |
| Pyruvate | 43.28 | **3.18E-08** | 7.11 | 0.49 | 40.44 | **8.76E-08** |
| Melibiose | 41.88 | **4.42E-08** | 2.64 | 0.93 | 36.48 | **3.84E-07** |
| Fumarate | 17.12 | **1.83E-03** | 0.72 | 0.95 | 34.21 | **7.77E-07** |
| Malate | 40.58 | **6.15E-08** | 15.63 | **0.03** | 31.61 | **2.46E-06** |
| Gluconate | 45.31 | **2.37E-08** | 23.54 | **0.001** | 47.69 | **8.55E-09** |
| Mannose | 39.93 | **7.45E-08** | 3.94 | 0.78 | 34.50 | **7.36E-07** |
| NAG | 38.76 | **1.17E-07** | 8.16 | 0.43 | 35.75 | **4.89E-07** |
| Glucuronate | 36.71 | **2.82E-07** | 1.58 | 0.94 | 47.61 | **8.55E-09** |

**Table S7. Output of two-sided Student's t-tests for differences between fitness of WT and mutator ancestor, in each environment.** p values are corrected for multiple comparisons using the Benjamini-Hochberg correction for false discovery rate.

| **Environment** | **Estimate** | **E1** | **E2** | **Statistic** | **p** | **Parameter** | **95% CI** | |
|---|---|---|---|---|---|---|---|---|
| | | | | | | | **Lower** | **Upper** |
| LB | -0.08 | 1.28 | 1.36 | -2.21 | 5.31E-02 | 80.02 | -0.154 | -0.008 |
| Glucose | 0.01 | 0.78 | 0.76 | 0.85 | 4.54E-01 | 85.23 | -0.017 | 0.043 |
| Trehalose | 0.01 | 0.49 | 0.48 | 1.08 | 3.51E-01 | 73.68 | -0.009 | 0.029 |
| Fructose | 0.03 | 0.57 | 0.54 | 2.95 | **1.39E-02** | 66.04 | 0.010 | 0.051 |
| Lactose | 0.02 | 0.75 | 0.73 | 1.65 | 1.38E-01 | 75.50 | -0.004 | 0.044 |
| Maltose | -0.02 | 0.52 | 0.54 | -3.06 | **1.39E-02** | 43.58 | -0.036 | -0.008 |
| Galactose | 0.04 | 0.35 | 0.31 | 5.18 | **2.67E-05** | 78.62 | 0.027 | 0.061 |
| Succinate | 0.02 | 0.49 | 0.47 | 1.94 | 9.65E-02 | 36.86 | -0.001 | 0.037 |
| Pyruvate | -0.02 | 0.44 | 0.46 | -2.77 | **2.03E-02** | 56.38 | -0.043 | -0.007 |
| Melibiose | 0.04 | 0.55 | 0.51 | 4.84 | **5.00E-05** | 79.52 | 0.022 | 0.052 |
| Fumarate | -0.03 | 0.42 | 0.45 | -5.01 | **5.48E-05** | 42.18 | -0.040 | -0.017 |
| Malate | 0.01 | 0.46 | 0.46 | 0.76 | 4.79E-01 | 71.86 | -0.009 | 0.020 |
| Gluconate | -0.06 | 0.61 | 0.67 | -1.91 | 1.15E-01 | 12.70 | -0.134 | 0.008 |
| Mannose | 0.03 | 0.30 | 0.27 | 2.62 | 5.00E-02 | 10.29 | 0.005 | 0.058 |
| NAG | -0.02 | 0.64 | 0.65 | -0.56 | 5.89E-01 | 8.76 | -0.079 | 0.048 |
| Glucuronate | 0.03 | 0.62 | 0.59 | 2.53 | 5.00E-02 | 13.83 | 0.005 | 0.056 |

**Table S8. Output of paired Wilcoxon's rank-sum tests to compare fitness effects of identical mutations across WT and mutator genetic backgrounds.**

| **Environment** | **Statistic** | **p** |
|---|---|---|
| LB | 117 | 0.839 |
| Glucose | 89 | 0.937 |
| Trehalose | 90 | 0.937 |
| Fructose | 114 | 0.839 |
| Lactose | 104 | 0.937 |
| Maltose | 102 | 0.937 |
| Galactose | 71 | 0.839 |
| Succinate | 151 | 0.194 |
| Pyruvate | 148 | 0.194 |
| Melibiose | 113 | 0.839 |
| Fumarate | 94 | 0.984 |
| Malate | 129 | 0.727 |
| Gluconate | 183 | **0.001** |
| Mannose | 86 | 0.916 |
| NAG | 118 | 0.783 |
| Glucuronate | 152 | 0.122 |

**Table S9. Output of two-sided ANOVAs to test the effects of mutation class and environment on fitness in WT, in each of three mutation bias categories where mutation spectra differ significantly between WT and mutator.**

| Mutation bias type | Term | df | Sum-sq. | Mean-sq. | Statistic | p |
|---|---|---|---|---|---|---|
| WT: Ts/Tv | Mutation class | 1 | 0.3962 | 0.3962 | 8.2265 | **0.0042** |
| | Environment | 15 | 0.7015 | 0.0468 | 0.9710 | 0.4839 |
| | Mutation class x Environment | 15 | 0.4519 | 0.0301 | 0.6255 | 0.8558 |
| | Residuals | 1120 | 53.942 | 0.0482 | NA | NA |
| WT: GC→AT | Mutation class | 1 | 0.6871 | 0.6871 | 13.234 | **0.0003** |
| | Environment | 15 | 0.5915 | 0.0394 | 0.7596 | 0.7235 |
| | Mutation class x Environment | 15 | 0.2053 | 0.0137 | 0.2637 | 0.9978 |
| | Residuals | 1008 | 52.330 | 0.0519 | NA | NA |
| WT: Coding/ Non-coding | Mutation class | 1 | 0.0924 | 0.0924 | 1.7334 | 0.1882 |
| | Environment | 15 | 1.0911 | 0.0727 | 1.3650 | 0.1566 |
| | Mutation class x Environment | 15 | 0.2026 | 0.0135 | 0.2534 | 0.9983 |
| | Residuals | 1248 | 66.506 | 0.0533 | NA | NA |
| WT: Synonymous/ Nonsynonymous | Mutation class | 1 | 0.1240 | 0.1240 | 2.8884 | 0.0896 |
| | Environment | 15 | 0.5419 | 0.0361 | 0.8413 | 0.6314 |
| | Mutation class x Environment | 15 | 0.0990 | 0.0066 | 0.1536 | 0.9999 |
| | Residuals | 864 | 37.103 | 0.0429 | NA | NA |
| Mutator: Synonymous/ Nonsynonymous | Mutation class | 1 | 0.0546 | 0.0546 | 4.3690 | **0.03684** |
| | Environment | 15 | 0.996 | 0.0664 | 5.3139 | **2.19E-10** |
| | Mutation class x Environment | 15 | 0.1520 | 0.0101 | 0.8107 | 0.666465 |
| | Residuals | 1040 | 13.007 | 0.0125 | NA | NA |

**Table S10. Output of two-sample Kolmogorov-Smirnov tests comparing the distributions of distances from *oriC* of synonymous, nonsynonymous and non-coding mutations; and distributions of distances from the start codon for synonymous and nonsynonymous coding mutations; between WT and mutator.**

| | Statistic | p |
|---|---|---|
| All mutations | | |
| Synonymous | 0.365 | 0.179 |
| Non-synonymous | 0.271 | 0.067 |
| Non-coding | 0.167 | 0.996 |
| Coding mutations | | |
| Synonymous | 0.161 | 0.986 |
| Non-synonymous | 0.117 | 0.881 |

**Table S11. Chi-square tests comparing the proportion of mutations with various characteristics, in WT vs. mutator.** In each category, we separately tested synonymous and non-synonymous mutations.

| Comparison | Mutation type | Statistic | p | Parameter |
|---|---|---|---|---|
| Replichore | Synonymous | 0.001 | 0.964 | 1 |
| | Non-synonymous | 0.074 | 0.785 | 1 |
| | Non-coding | 0.823 | 0.364 | 1 |
| Strand | Synonymous | 0.086 | 0.770 | 1 |
| | Non-synonymous | 2.432 | 0.119 | 1 |
| Core vs. Accessory gene | Synonymous | 0.000 | 1.000 | 1 |
| | Non-synonymous | 0.003 | 0.955 | 1 |
| Essentiality in LB | Synonymous | 0.161 | 0.688 | 1 |
| | Non-synonymous | 0.107 | 0.743 | 1 |
| Essentiality in M9 minimal medium | Synonymous | 0.878 | 0.644 | 2 |
| | Non-synonymous | 1.465 | 0.481 | 2 |
| GO category | Synonymous | 4.042 | 0.671 | 6 |
| | Non-synonymous | 3.104 | 0.683 | 6 |
| Amino-acid change type | Non-synonymous | 0.028 | 0.867 | 1 |

**Table S12. Experimentally validated effects of DNA repair gene loss on mutation biases in *Escherichia coli* strains.** Relative bias values of 1 indicate the same absolute bias as WT *E. coli*; values > 1 indicate more Tv or GC→AT mutations than WT *E. coli*; values <1 indicate more Ts or AT→GC mutations than WT *E. coli.*

| Strain | Mutation rate ($bp^{-1}$ $gen^{-1}$)[a] | Absolute bias[b] | | Relative bias[c] | | Reference |
|---|---|---|---|---|---|---|
| | | Tv bias | GC→AT bias | Tv bias | GC→AT bias | |
| WT[$] | 2.05E-10 | 0.447 | 0.557 | 1.000 | 1.000 | |
| uvrA | 2.28E-10 | 0.472 | 0.558 | 1.055 | 1.003 | |
| nfi | 2.44E-10 | 0.457 | 0.533 | 1.022 | 0.957 | (4) |
| mutT | 3.22E-08 | 0.997 | 0.002 | 2.231 | 0.004 | |
| mutY | 2.21E-09 | 0.948 | 0.956 | 2.122 | 1.717 | |
| dnaQ* | 8.44E-07 | 0.140 | 0.439 | 0.313 | 0.787 | (31) |
| mutL** | 2.46E-08 | 0.032 | 0.232 | 0.072 | 0.416 | (32, 33) |
| mutS** | 2.76E-08 | 0.025 | 0.205 | 0.055 | 0.368 | |
| mutH | 2.35E-08 | 0.029 | 0.294 | 0.066 | 0.528 | (33) |
| uvrD | 1.29E-08 | 0.043 | 0.338 | 0.096 | 0.607 | |
| ung | 3.84E-10 | 0.159 | 0.832 | 0.356 | 1.494 | This study |
| mutM | 2.57E-10 | 0.563 | 0.692 | 1.258 | 1.243 | |
| mutL mutY | 2.89E-08 | 0.081 | 0.238 | 0.182 | 0.427 | |
| mutL mfd | 2.34E-08 | 0.024 | 0.181 | 0.053 | 0.326 | |
| mutS mfd | 3.02E-08 | 0.025 | 0.209 | 0.055 | 0.376 | (33) |
| mutL ndk | 1.32E-07 | 0.049 | 0.048 | 0.109 | 0.086 | |
| mutL mutS | 2.43E-08 | 0.030 | 0.164 | 0.066 | 0.295 | |
| alkA tagA | 1.95E-10 | 0.445 | 0.563 | 0.996 | 1.011 | |
| umuDC dinB | 2.22E-10 | 0.494 | 0.634 | 1.106 | 1.138 | |
| ada ogt | 1.75E-10 | 0.432 | 0.632 | 0.966 | 1.134 | |
| nth nei | 9.79E-10 | 0.493 | 0.375 | 1.103 | 0.673 | (4) |
| xthA nfo | 5.26E-10 | 0.499 | 0.516 | 1.115 | 0.927 | |
| mutM mutY | 2.01E-08 | 0.994 | 0.996 | 2.225 | 1.788 | |
| dnaQ* mutL | 1.25E-06 | 0.062 | 0.632 | 0.138 | 1.134 | |
| dnaQ* dinB | 7.19E-07 | 0.120 | 0.462 | 0.268 | 0.830 | (31) |
| dnaQ* lexA3 | 1.16E-06 | 0.102 | 0.497 | 0.228 | 0.892 | |
| umuDC dinB polB | 1.95E-10 | 0.437 | 0.610 | 0.977 | 1.096 | (4) |
| mutL mutS mutH[#] | 2.10E-08 | 0.023 | 0.307 | 0.051 | 0.551 | (33) |
| mutL umuDC dinB | 1.98E-08 | 0.027 | 0.264 | 0.060 | 0.474 | |
| dnaQ* dinB umuDC | 9.01E-07 | 0.125 | 0.478 | 0.281 | 0.858 | (31) |

[a] Mutation rate was calculated as: Total number of base-pair substitutions/(Genome size x number of generations per lineage x total number of lineages)

[b] Absolute bias was calculated as: Tv bias = Number of transversions/(Number of transversions + transitions); GC→AT bias = Number of GC→AT mutations/(Number of GC→AT mutations + AT→GC mutations)

[C] Relative biases were calculated by dividing the absolute bias of each strain by the absolute bias of WT *E. coli*

[$] WT *E. coli* reference

* Data are for the mutD5 allele of dnaQ, which reduces exonuclease activity by 98% but retains core polymerase structure (17)

** Average values from 3 independent sets of MA experiments

[#] Average values from 2 independent sets of MA experiments

**Table S13. Impact of loss of DNA repair genes across different species.** Absolute biases were calculated as in Table S12. Relative biases were calculated as the ratio of the absolute bias values of the deletion strain, divided by the respective wild type strain from the same study.

| Organism | Gene deleted or inactivated | Absolute bias | | Relative bias | | Reference |
|---|---|---|---|---|---|---|
| | | Tv bias | GC→AT bias | Tv bias | GC→AT bias | |
| *Escherichia coli* | mutS | 0.02 | 0.20 | 0.05 | 0.4 | (33) |
| *Bacillus subtilis* | mutS | 0.03 | 0.51 | 0.11 | 1.02 | (19) |
| *Vibrio fischeri* | mutS | 0.02 | 0.40 | 0.04 | 0.55 | |
| *Vibrio cholerae* | mutS | 0.02 | 0.50 | 0.06 | 0.80 | (34) |
| *Pseudomonas fluorescens* | mutS | 0.01 | 0.49 | 0.03 | 0.76 | (35) |
| *Pseudomonas aeruginosa* | mutS | 0.01 | 0.52 | 0.03 | 0.70 | (23) |
| *Deinococcus radiodurans* | mutS1[a] | 0.06 | 0.47 | 0.16 | 0.90 | (22) |
| *Escherichia coli* | mutL | 0.033 | 0.23 | 0.07 | 0.42 | (4, 32) |
| *Vibrio cholerae* | mutL | 0.01 | 0.46 | 0.03 | 0.76 | (21) |
| *Pseudomonas fluorescens* | mutL | 0.01 | 0.50 | 0.03 | 0.77 | (35) |
| *Deinococcus radiodurans* | mutL | 0.05 | 0.64 | 0.13 | 1.22 | (24) |

[a] One of the two mutS homologs in *Deinococcus radiodurans*. Deletion of mutS1, but not mutS2, causes increased mutation rate (36).

[b] msh2 is a mutS homolog in *Saccharomyces cerevisiae*, involved in mismatch repair in the nucleus (37).
